# Sources of Off-Target Effects of Vagus Nerve Stimulation Using the Helical Clinical Lead in Domestic Pigs

**DOI:** 10.1101/2020.01.15.907246

**Authors:** Evan N. Nicolai, Megan L. Settell, Bruce E. Knudsen, Andrea L. McConico, Brian A. Gosink, James K. Trevathan, Ian W. Baumgart, Erika K. Ross, Nicole A. Pelot, Warren M. Grill, Kenneth J. Gustafson, Andrew J. Shoffstall, Justin C. Williams, Kip A. Ludwig

## Abstract

Clinical data suggest that efficacious vagus nerve stimulation (VNS) is limited by side effects such as cough and dyspnea that have stimulation thresholds lower than those for therapeutic outcomes. VNS side effects are putatively caused by activation of nearby muscles within the neck, via direct muscle activation or activation of nerve fibers innervating those muscles. Our goal was to determine the thresholds at which various VNS-evoked effects occur in the domestic pig—an animal model with vagus anatomy similar to human—using the bipolar helical lead deployed clinically. Intrafascicular electrodes were placed within the vagus nerve to record electroneurographic (ENG) responses, and needle electrodes were placed in the vagal-innervated neck muscles to record electromyographic (EMG) responses. Contraction of the cricoarytenoid muscle occurred at low amplitudes (∼0.3 mA) and resulted from activation of motor nerve fibers in the cervical vagus trunk within the electrode cuff which bifurcate into the recurrent laryngeal branch of the vagus. At higher amplitudes (∼1.4 mA), contraction of the cricoarytenoid and cricothyroid muscles was generated by current leakage outside the cuff to activate motor nerve fibers running within the nearby superior laryngeal branch of the vagus. Activation of these muscles generated artifacts in the ENG recordings that may be mistaken for compound action potentials representing slowly conducting Aδ-, B-, and C-fibers. Our data resolve conflicting reports of the stimulation amplitudes required for C-fiber activation in large animal studies (>10 mA) and human studies (<250 µA). After removing muscle-generated artifacts, ENG signals with post-stimulus latencies consistent with Aδ- and B-fibers occurred in only a small subset of animals, and these signals had similar thresholds to those that caused bradycardia. By identifying specific neuroanatomical pathways that cause off-target effects and characterizing the stimulation dose-response curves for on- and off-target effects, we hope to guide interpretation and optimization of clinical VNS.

## Introduction

The cervical vagus nerve provides an entry point for modulating both visceral organ function and much of the brain. The human cervical vagus contains a complex mixture of sensory afferents that project to the nucleus tractus solitarius, which projects to diffuse regions of the brain (Altschuler et al., 1989; Kalia & Mesulam, 1980), preganglionic parasympathetic efferents projecting to the visceral organs (Agostoni et al., 1957; Foley & DuBois, 1937; Hoffman & Schnitzlein, 1961; Mei et al., 1980), and somatic motor efferents projecting to muscles of the neck. Therapeutic vagus nerve stimulation (VNS) with implanted electrodes has grown over recent years for diverse conditions from epilepsy to heart failure (De Ferrari et al., 2017; Kimberley et al., 2018; Koopman et al., 2016; Morris et al., 2013; Ng et al., 2016; Tyler et al., 2017; Wheless et al., 2018). Despite remarkable outcomes in some patients across numerous pathologies overall results are often mixed, with side effects reported in up to 66% of patients that include cough, throat pain, voice alteration, difficulty swallowing, and dyspnea (De Ferrari et al., 2017; A. Handforth et al., 1998; VNS Study Group, 1995). These side effects can limit tolerable stimulation amplitudes to below effective therapeutic levels. In an early study of VNS for epilepsy in humans, patients were unable to tolerate stimulation amplitudes higher than 1.3 mA on average using a 500 µs pulse width at 30 Hz (A. Handforth et al., 1998). Similarly the average tolerable amplitude was 1.2 mA using a 300 µs pulse width at 20 Hz in a recent clinical trial, with only 12% of patients experienced targeted VNS-evoked heart rate changes 1 year post-implant (De Ferrari et al., 2017).

To guide efforts to minimize VNS therapy limiting side effects, we isolated the neuroanatomical pathways responsible for VNS-induced neck muscle contraction using the FDA approved bipolar helical clinical lead from LivaNova. We also characterized the dose response curves for unwanted neck muscle activation in comparison to activation of nerve fiber types within the cervical vagus and stimulation induced changes in heart rate. We hypothesized that activation of efferent A-type motor nerve fibers within or near the vagus nerve were causing neck muscle contraction in response to VNS, as opposed to direct electrical activation of the muscles. As the vagus has multiple somatic branches, we hypothesized that cervical VNS would 1) activate motor fibers in the cervical vagus trunk within the cuff electrodes that eventually bifurcate into the recurrent laryngeal branch of the vagus at low amplitudes, and 2) activate the motor fibers in the nearby superior laryngeal branch of the vagus ‘en passant’ at higher amplitudes (Settell et al., 2019).

To test these hypotheses, anesthetized domestic pigs were acutely stimulated with the bipolar helical lead used for clinical VNS in an intact state, under neuromuscular junction block, and after dissections of the recurrent and superior laryngeal branches. The domestic pig was chosen due to its anatomical similarities to human (Ding et al., 2012; Settell et al., 2019). Electromyography (EMG) recordings of the terminal neck muscles of the recurrent/superior laryngeal branches under each condition were obtained. Simultaneously, longitudinal intrafascicular electrode (LIFE) recordings were obtained from electrodes sewn into the cervical vagus nerve. Dose response curves for EMG and LIFE electrode recordings during VNS were directly compared to dose response curves for stimulation induced reductions in heart rate (bradycardia).

By identifying the neuroanatomical pathways responsible for off-target effects, we hope these data can guide new strategies to avoid these unwanted effects to maximize intended therapeutic effect. We also provide the first comprehensive dataset in an animal model with the most similar vagus nerve size and anatomy to humans (Settell et al., 2019) using an unmodified and unscaled helical clinical lead.

## Methods

All animal care and procedures were approved by the Institutional Animal Care and Use Committee of Mayo Clinic. Twelve domestic pigs were included in the study: seven male pigs, weighing 37.7 ± 2.3 kg (mean ± standard deviation, SD) and five female pigs weighing 38.3 ± 3.9 kg. All procedures described were performed on either the left or right vagus nerve, and only one side per animal. All procedures were performed in an acute terminal study lasting between 10 and 12 hours.

### Anesthesia and Surgical Procedure

All animals were sedated using an intramuscular injection of telazol (6 mg/kg), xylazine (2 mg/kg), and glycopyrrolate (0.006 mg/kg), then intubated and anesthetized using isoflurane (1.5-3% isoflurane in room air). Saline (0.9%) was administered continuously with analgesia (fentanyl, 5 µg/kg bolus i.v., followed by 5 µg/kg/hr in saline drip). Heart rate, respiration, blood pressure, and end-tidal CO2 were used to assess depth of anesthesia, and isoflurane dosage was adjusted as needed. Animals were euthanized using pentobarbital (100 mg/kg i.v.).

Pigs were positioned in supine position with neck extended, and a single incision was made through the skin and superficial fat layers between the mandible and sternal notch using a cautery. Blunt dissection was used to expose the carotid sheath and to isolate the vagus nerve from the carotid artery. Care was taken to avoid perturbation of the vagus nerve, while other structures were pulled away from the vagus nerve using vessel loops. The surgical window was kept open throughout the procedure. Pooled liquid in the surgical window was removed periodically, and saline was sprayed evenly through the window to keep all structures moist.

### Electrode Types and Surgical Placement

The placement of electrodes in a typical surgical window is shown in Figure 1. The clinical bipolar stimulating electrode (PerreniaFLEX Model 304, 2 mm inner diameter, LivaNova, Houston, TX) was purchased and used without modification. A practicing neurosurgeon assisted in choosing a placement location most similar to human patients. The two contacts were always placed caudal to the superior laryngeal branching point.

**Figure 1:**
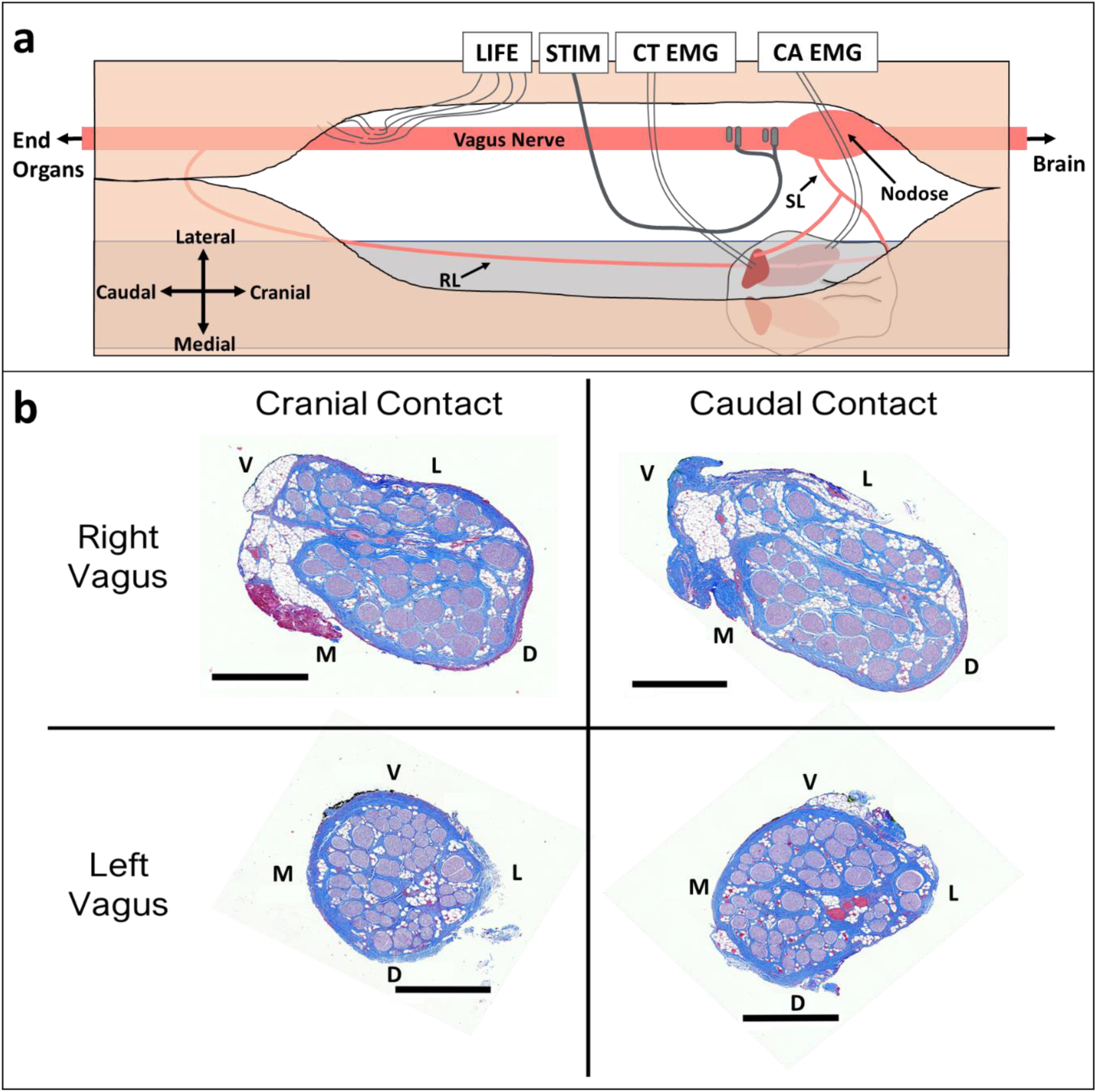
Vagus nerve anatomy and experimental setup. **a)** Diagram (not to scale) of surgical window showing key anatomy and relative electrode locations. Longitudinal intrafascicular electrode (LIFE), bipolar helical clinical stimulation lead (STIM), cricothyroid muscle (CT), cricoarytenoid muscle (CA), electromyography (EMG), superior laryngeal (SL), recurrent laryngeal (RL). **b)** Representative cross sections from left and right vagus nerves obtained from the location along the vagus nerve where stimulation electrode contacts were placed. Black scale bars are 1 mm. Letters indicate anatomical directions ventral (V), lateral (L), medial (M), and dorsal (D).

Bipolar stainless steel needle electrodes were used for measurement of electromyograms (EMG) in the cricothyroid and cricoarytenoid muscles. Custom longitudinal intrafascicular electrodes (LIFEs) were built in-house for measuring electroneurograms (ENG) (based on Lefurge et al., 1991) taken within the cervical vagus. LIFE electrode recordings were used over epineural cuff recordings, as a prior study of VNS demonstrated that recording activity from low amplitude C-fibers is difficult without shaving the epineurium to reduce the distance from the C-fiber signal source to the epineural electrode (Yoo et al., 2013). Briefly, to fabricate LIFE electrodes, 300 to 500 µm of insulation was removed at a single location along a 75 µm outer diameter PFA-coated platinum wire (AM-Systems, Sequim, WA) using a thermal wire stripper to create an electrode contact. A suture tie was placed near the location of the electrode contact to assist in tracking the electrode contact during the surgical procedure. A suture needle (Item No. 12050-03, Fine Science Tools, Foster City, CA) was attached to one end of the platinum wire by threading the wire through the needle eyelet and twisting the wire, and an insulated copper extension wire with touchproof connector (441 connector with wire, Plastics1, Roanoke, VA) was soldered to the other end. The needle was used to introduce each LIFE into the vagus nerve, and the needle was removed prior to recordings. Three to five LIFEs were implanted per animal in a staggered cluster (Figure 1a) where the center LIFE was 8.6 ± 1.1 cm caudal to the location of the center of the cranial stimulation electrode contact. Placement of the LIFEs was staggered to prevent exact matching of ENG signal latencies, which could cause the target neural signal to be lost during differential subtraction.

### Equipment

All LIFE and EMG recordings, as well as stimulations, were performed using a Tucker Davis Technologies system (Alachua, FL; W8, IZ2MH, RZ5D, RZ6, PZ5, and S-Box units). These recordings were collected continuously (24,414 Hz sampling rate, 10,986 Hz anti-aliasing, unity gain) before, during, and after application of each stimulation pulse. One of the LIFEs was used as the reference electrode for all electrophysiology recordings.

Heart rate (HR) and blood pressure were monitored using a pressure catheter (Millar Inc, Houston, TX, Model #SPR-350S) placed into the femoral artery, then digitized and saved using a PowerLab and Bridge Amplifier (ADInstruments, Sydney, Australia).

### Experimental Protocol

Bipolar stimulation was delivered at 30 Hz using symmetric biphasic pulses with 200 µs per phase and amplitudes from 50 to 3000 µA evaluated in a random order. Stimulation was typically delivered for 30 seconds at each amplitude with at least 1 min rest between trials to allow cardiac responses to return to baseline; in some cases, stimulation was delivered for 3 seconds at each amplitude with at least 10 seconds rest due to time constraints. Stimulation parameters were chosen to approximate the parameters used in the clinic (De Ferrari et al., 2017; A. Handforth et al., 1998; VNS Study Group, 1995). In the default bipolar configuration, the more cranial electrode delivered the cathodic phase first. In a subset of animals, stimulation cathode and anode were reversed.

Input-output curves were generated for ENG, EMG, and HR from 50 to 3000 µA (including at least 100, 200, 300, 400, 500, 750, 1000, 1500, 2000, 2500, and 3000 µA presented) for both cathode cranial and cathode caudal configurations. To verify the source of EMG and ENG recordings, some stimulation amplitudes – typically 1000, 2000, and 3000 µA – were repeated following a series of perturbations including administration of a neuromuscular blocking agent (vecuronium, 0.1 mg/kg bolus i.v., followed by 3 mg/kg/hr continuous), transection of the recurrent and superior laryngeal vagus nerve branches, and transection of the vagus trunk between the stimulation electrode and LIFEs.

### Data Analysis

All signal processing was performed in Matlab R2018b (Mathworks, Natick, MA). ENG and EMG recordings were filtered using a 400 sample median high-pass filter (y=x-medfilt1(x)) and a gaussian low-pass filter (gaussfiltcoef and filtfilt) with corner frequency of 5 kHz; these filters were applied in order to lessen filter ringing near the stimulation artifact that may obscure recording of the fastest compound action potentials. Stimulation-triggered median traces of evoked LIFE and EMG recordings were calculated for each stimulation amplitude. The activation threshold and latency for each feature were visually identified in the compound action potential and EMG traces (Figure 2). For LIFE recordings, conduction velocities and measured distances between the electrode delivering cathode phase first and a given LIFE were used to determine latencies for each fiber type (Aα 70-120 m/s, Aβ 40-70 m/s, Aγ 15-40 m/s, Aδ 5-15 m/s, B 3-14 m/s) (Erlanger & Gasser, 1937; Manzano et al., 2008; Parent & Carpenter, 1996). Aα-fibers are not reported here since the short expected latencies (<1 ms) of these fibers were coincident with the stimulation artifact and could not be directly measured. Aα-fibers are larger diameter than Aβ-fibers, and thus have lower stimulation thresholds; therefore, we made the assumption that measurement of ENGs with post-stimulus latencies consistent with Aβ-fibers was an indication of Aα-fiber activation at a lower threshold. Aδ- and B-fibers have overlapping conduction velocities and were therefore treated as one combined neural response (Aδ/B). C-fiber components were not evident in any of our recordings with one possible exception (discussed later). For most animals, two EMG responses occurred with distinct stimulation thresholds and distinct latencies. The peak of the first major deflection was used to calculate the latency for each response. Data is presented comparing the left and right vagus responses, as well as between male and female animals, but no statistical comparisons were made due to relatively low number of animals in each group (12 total pigs, 5 left side vs 7 right side, 5 female vs 7 male). A summary of all data points is available in Figure 6 and Supplementary Figure 4.

**Figure 2:**
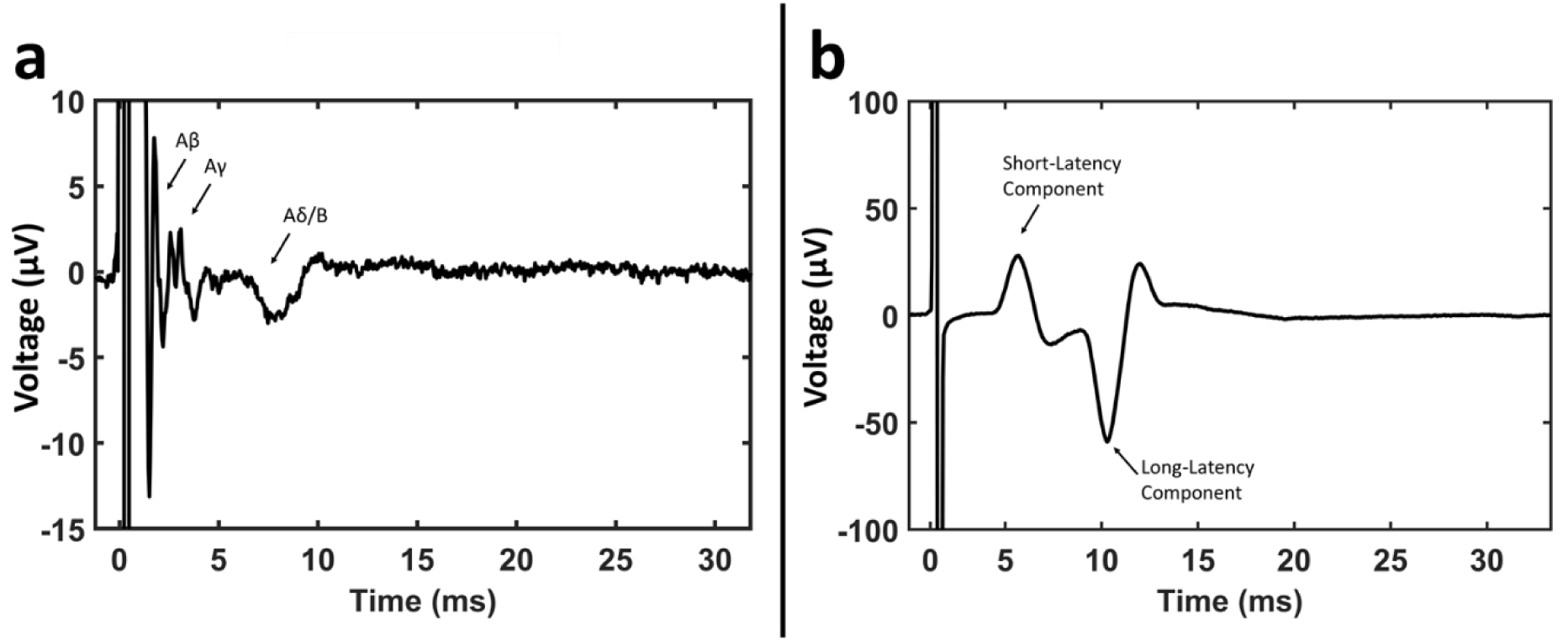
Example ENG and EMG recordings and classification. **a)** Example recording from a LIFE showing compound action potential features. **b)** Example EMG recording in the cricothyroid muscle showing short- and long-latency components, with the first major deflection of each component labeled, used to calculate the latencies.

**Figure 3:**
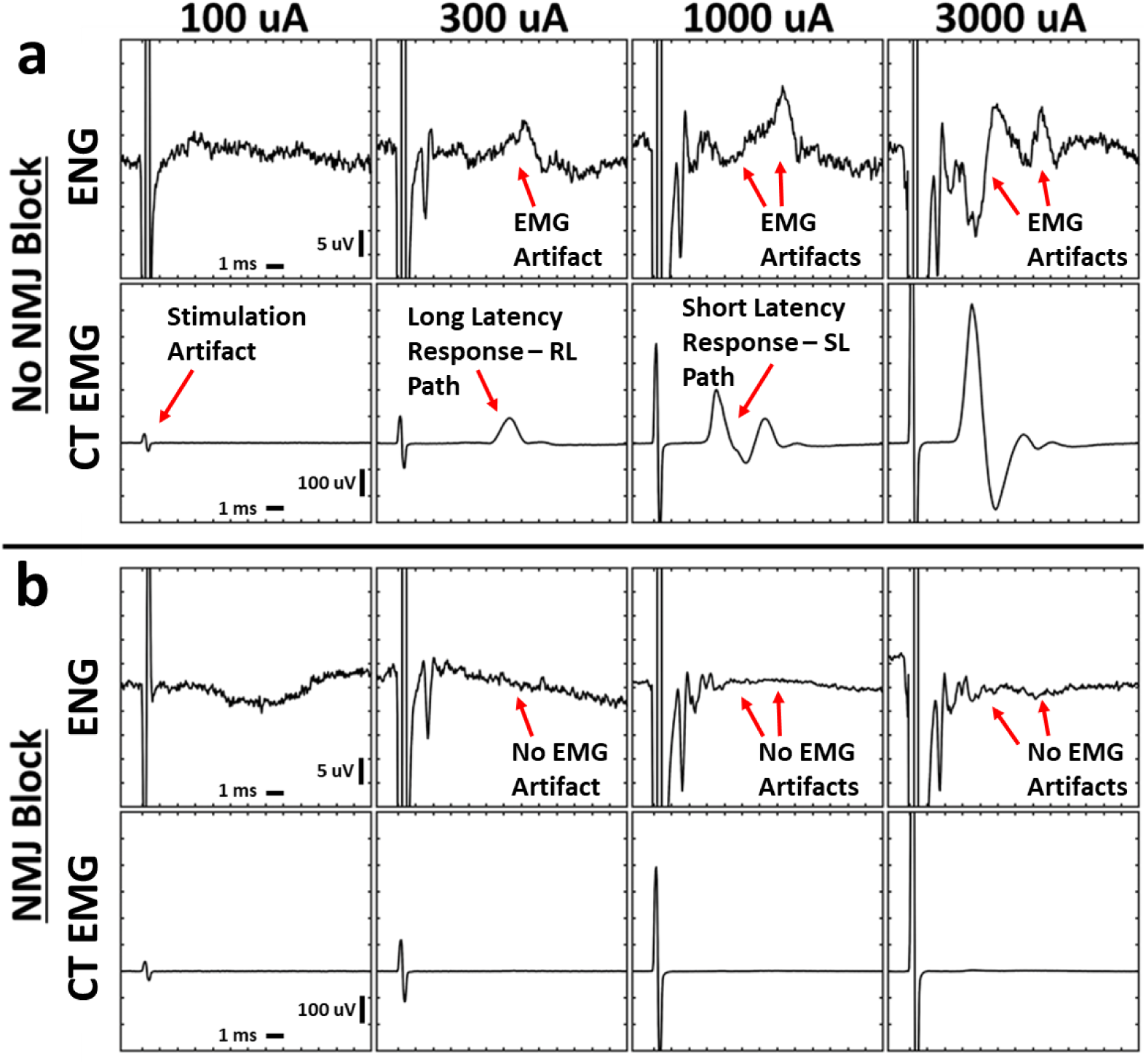
Stimulation-triggered median ENG and EMG evoked by cervical VNS before and after neuromuscular junction blockade. Electromyograms (EMG) exhibited short- and long-latency components at distinct thresholds, and these signals contaminated the electroneurograms (ENG). Neuromuscular blockade with vecuronium eliminated all cricothyroid (CT) and cricoarytenoid (CA, not shown) EMGs and EMG contamination of ENGs. **a)** Simultaneously collected ENG and EMG at multiple stimulation amplitudes (columns) without neuromuscular blockade. **b)** Analogous data in the same animal following neuromuscular blockade. X-axis ticks in all plots are 1 ms. Y-axis ticks are 5 µV in all ENG plots and 100 µV in all EMG plots.

**Figure 4:**
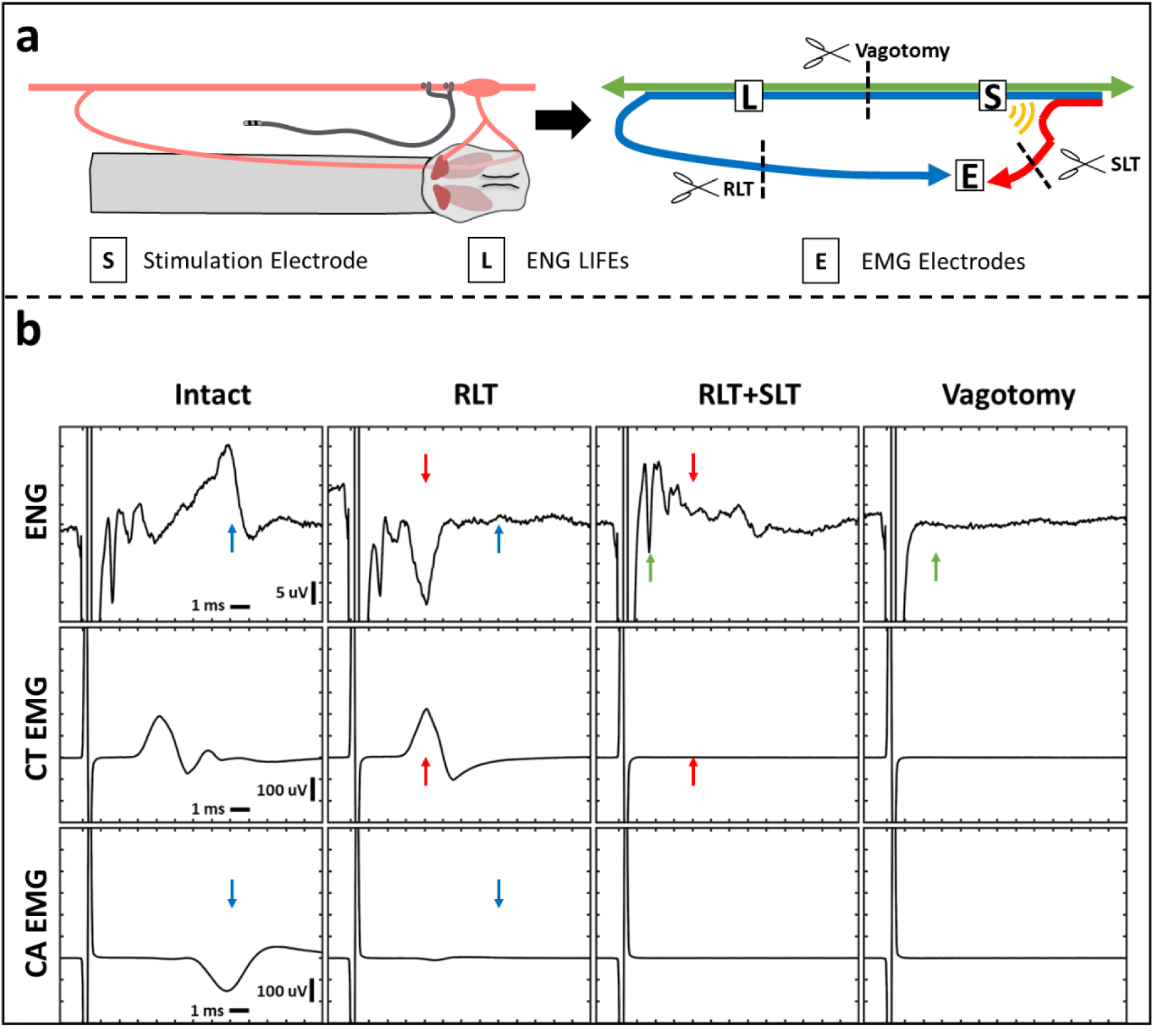
ENG and EMG Recordings before and after Transections. Transection of recurrent laryngeal branch (RLT) and superior laryngeal branch (SLT) eliminated long- and short-latency EMGs, respectively. Transection of the vagus trunk eliminated all ENGs. **a)** Diagram of the transection methods. Left panel shows a diagram of the surgical window. Right panel shows a wiring diagram of nerve fibers with transection locations (scissors). Blue line depicts recurrent laryngeal branch fibers, red line depicts superior laryngeal branch fibers, and green line depicts other vagus nerve fibers. Yellow semi-circles indicate expected current leakage out of the nerve cuff insulation acting on the superior laryngeal branch fibers that lie outside the cuff. **b)** One channel of ENG (top row) and two channels of EMG (cricothyroid (CT; middle row) and cricoarytenoid (CA; bottom row)) collected simultaneously. Columns represent different time points in order from left to right with an additional transection at each step starting from no transections (Intact, first column). Stimulation parameters for every column are charge-balanced symmetrical biphasic pulses at 3 mA with 200 µs per phase, delivered at 30 Hz for 30 seconds. Paired colored arrows highlight components of the signal that changed before and after each transection.

**Figure 5:**
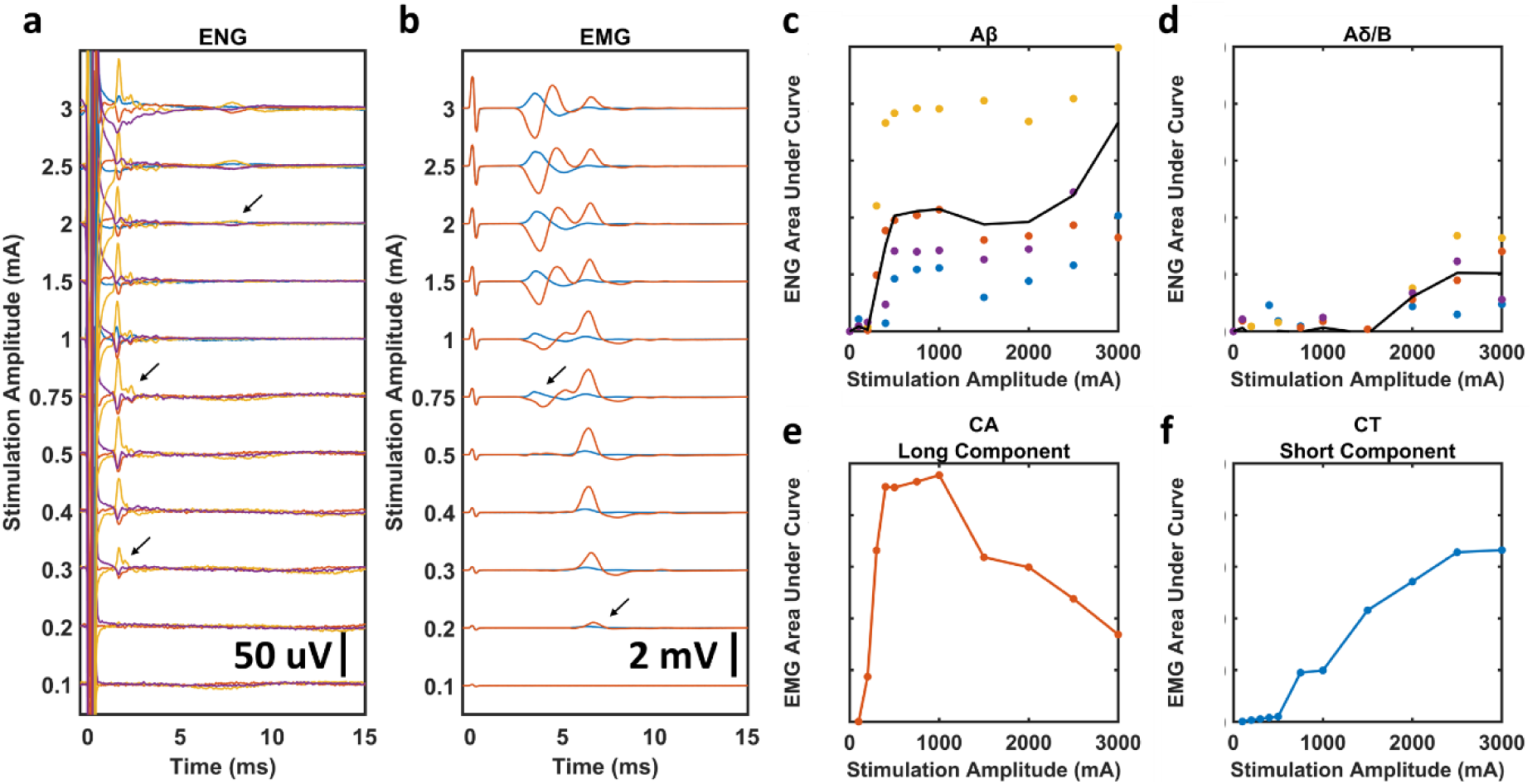
Stimulation Dose-Response Curves and Determination of Stimulation Amplitude Threshold for each Fiber Type in a Representative Animal. EMG recordings were taken before neuromuscular blockade, ENG recordings were taken in the same animal during neuromuscular blockade. **a)** All stimulation-triggered median ENGs for a representative animal. Arrows indicate visually identified thresholds for each ENG signal; Aβ, Aγ, and Aδ/B from bottom to top. Colored traces represent four different LIFEs. **b)** All stimulation-triggered median EMGs for the same representative animal. Arrows indicate visually identified thresholds for each EMG signal; long-component and short-component from bottom to top. **c-d)** Dose-response curves for Aβ and Aδ/B ENGs calculated using historically available conduction velocities to determine latency ranges. Black line is average of all LIFEs. **e-f)** Dose-response curves for long-component CA and short-component CT EMGs calculated using visually identified latency ranges.

**Figure 6:**
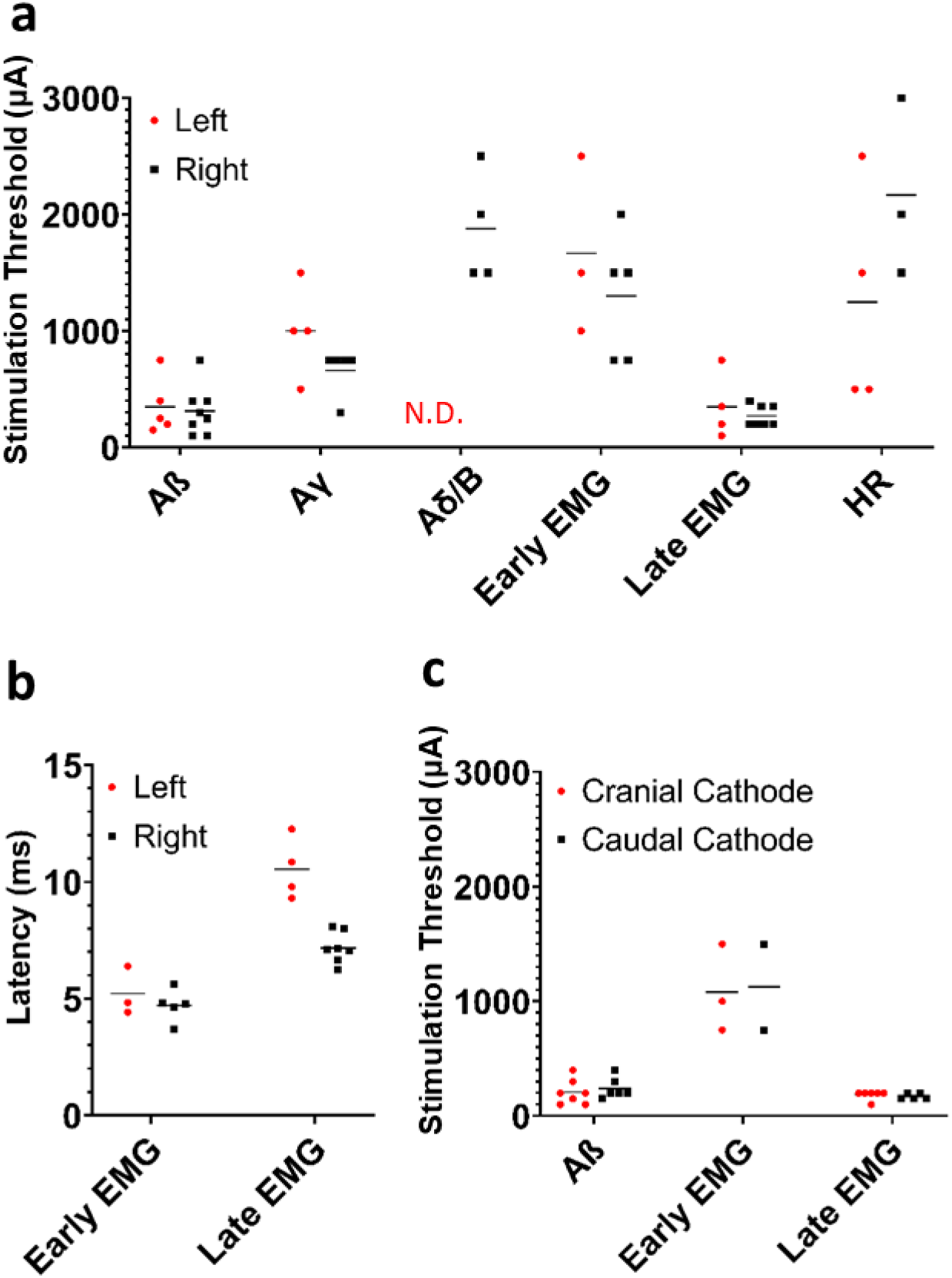
Summary of Comparisons Between Left and Right Side VNS, as well as Cathode and Anode configurations. **a)** Thresholds for ENG, EMG, and HR responses comparing left and right vagus nerve experiments. **b)** Post-stimulus latencies for EMG components comparing left and right vagus nerve experiments. **c)** Thresholds for ENG and EMG responses comparing anode and cathode configurations. Note that only animals where responses to both cathode cranial and cathode caudal configurations were recorded are plotted for panel c.

Heart rate data was first smoothed (smooth Matlab function) by 1000 data points. Baseline was calculated as the median of the 30 seconds of data prior to stimulation. The change in heart rate was calculated as the median of the 2 data points before and after the peak deflection (5 data points) subtracted from the baseline. See Supplementary Figure 5 for an example heart rate deflection during stimulation. Heart rate was also monitored in real time during every experiment, and real time monitored changes were used for heart rate measurements in the case of excessive noise in digitally collected heart rate. Threshold for heart rate change was defined as the first stimulation amplitude that produced a 3 beat per minute change and subsequent recovery, where the next highest stimulation amplitude also produced at least a 3 beat per minute change.

The raw data, experiment notes, Matlab code used to process all data with documentation, and Matlab figure files can be found on the SPARC Data & Resource Center (RRID:SCR_017041).

### Histological Analysis

Histology dye (Bradley Products, Inc., Davidson Marking System, Bloomington, MN) was placed along the ventral and lateral edge of the vagus nerve to maintain orientation. The nerve was sampled from just cranial to the nodose ganglion to just caudal to the recurrent laryngeal bifurcation, thus including the region of nerve where the stimulating electrode contacts were placed. The samples were placed in 10% neutral buffered formalin for ∼24 hr at 4°C. Samples underwent a series of standard processing steps as previously described (Settell et al. 2020). Each sample was then embedded in paraffin wax and allowed to set. The block was placed into an ice water bath for approximately one hour to rehydrate the tissue. The block was cut into 5 μm sections using a Leica Biosystems Rotary Microtome (Buffalo Grove, Illinois) and stained using Gomori’s trichrome. Slides were imaged using a Motic Slide Scanner (Motic North America, Richmond, British Columbia) at 20x.

## Results

Two known principles guided the formulation of our hypotheses for the neuroanatomical pathways most likely to cause neck muscle activation during VNS. First, the thresholds for direct activation of denervated muscle fibers are much higher than for activation of large diameter myelinated motor nerve fibers innervating those muscles (Mortimer, 1981). Second, the electric field from an electrode source decreases rapidly with distance from the electrode (Plonsey & Barr, 1995). These principles combined suggest that the likely sources for off-target neck muscle activation are easily activated large diameter motor fibers either within the cervical vagus itself, or large diameter motor fibers located near the placement of the helical epineural cuff on the cervical vagus.

It is well known that a subset of motor fibers within the cervical vagus eventually bifurcate into the recurrent laryngeal branch (RL) of the vagus and terminate in the cricoarytenoid muscle of the neck. The motor fibers of the superior laryngeal branch (SL) bifurcate from the cervical vagus cranial to the standard VNS electrode placement to terminate in the cricothyroid and cricoarytenoid muscles in both humans and pigs (Settell et al., 2019). The distance between the placement of the center of the cranial contact and the superior laryngeal bifurcation was on average 0.49 ± 0.24 cm across the pig cohort, and the distance between the center of the caudal contact and the superior laryngeal bifurcation was 1.14 ± 0.21 cm. Consequently, the SL was the closest source of motor fibers that could potentially be activated by current leakage escaping the helical epinural cuff. Hence we hypothesized that the motor fibers within the cervical vagus that become the RL would be activated at very low thresholds, as these are located within the epineural cuff. At higher amplitudes, we hypothesized that current leakage escaping the insulation would activate the motor fibers of the nearby SL.

### Neuromuscular Junction Blockade to Identify Signals Caused by Muscle Contraction in ENG and EMG Recordings

Previous studies have shown that neural recordings of evoked compound action potentials (CAPs) taken using epineural cuffs placed on the vagus can be contaminated by the activation of nearby muscles. To assess this possibility for intrafascicular electrodes sewn into the cervical vagus, VNS dose response curves were obtained in each animal before and after the administration of neuromuscular junction blockade (See Figure 3). Direct ‘myogenic’ activation of muscle fibers caused by current leakage outside of the epineural cuff electrode should be instantaneous and therefore overlap temporally with the stimulus artifact in the EMG recordings. In contrast, activation of motor nerve fibers innervating muscle would result in a delay before the EMG response. This delay is the sum of the time it takes for the action potential generated at a specific point in the motor nerve to the neuromuscular junction with its terminal muscle, and the inherent delay at the neuromuscular junction before fiber activation (Katz & Miledi, 1965).

At low levels of stimulation, a long latency response was consistently evident in EMG recordings from the cricothyroid muscle across the cohort (8.4 ± 1.9 ms after the stimulus artifact, see Figure 3 top for a representative example, see supplementary table 1 for summary of measurements in all animals). This long-latency delay indicates that the initial cricothyroid muscle response was caused by the activation of motor nerve fibers that traversed a long distance prior to reaching their terminal muscle. Across the cohort, the average distance from the epineural cuff to the recurrent laryngeal branch summed with the average distance from the recurrent laryngeal branch to the cricothyroid muscle was 27.9 ± 6.8 cm. Erlanger and Gasser reported the velocity of motor Aα-fibers as 70-120 m/s. Based on these two pieces of information, action potentials in average Aα-fibers (95 m/s) produced under the stimulation electrode should reach the cricothyroid muscle via the recurrent laryngeal pathway in 2.22-3.65 ms. Katz and Miledi reported the inherent synaptic delay of the neuromuscular junction for ex vivo sartorius muscle to be 1-3 ms; in this cohort of in vivo pig experiments, the synaptic delay was found to be approximately 4.4 ms by correlating the latency of EMG responses to the length of the path between stimulation electrode and muscle for every animal (see Supplementary Figure 6). Summing these two delays, the range becomes 6.62-8.05 ms which overlaps the experimentally obtained range of 6.5-10.3 ms, thus supporting VNS-evoked muscle contraction via the vagus recurrent laryngeal pathway. A response with identical latency was also observed in the LIFE recordings obtained without neuromuscular junction blockade (Figure 3a top row) but was eliminated after administration of neuromuscular junction blockade confirming this response as EMG artifact (Figure 3b top row). This EMG artifact could easily be confused with an Aδ/B-fiber neural response based on latency if using the Erlanger/Gasser scale (Erlanger & Gasser, 1937).

At high levels of stimulation, a shorter latency response of 4.9 ± 0.8 ms was in EMG recordings. This shorter latency delay indicates that a new motor nerve fiber pathway was recruited that was shorter distance from the point of action potential initiation to terminal neck muscle. Across the cohort, the distance from the branching point of superior laryngeal to the cricothyroid muscle was 3.0 ± 0.5 cm. Using the same line of reasoning as the long latency EMG component, the action potentials produced in motor Aα-fibers within the superior laryngeal branch will reach the cricothyroid muscle in 0.26-0.37 ms. Summed with the neuromuscular junction delay of 4 ms, the expected latency would be 4.66-4.77 ms which again overlaps the experimentally obtained 4.1-5.7 ms latency. A response with identical short latency was also observed in the LIFE recordings obtained without neuromuscular junction blockade (Figure 3a top row), but was eliminated after administration of neuromuscular junction blockade, confirming this response as EMG artifact (Figure 3b top row). This shorter latency EMG artifact in the LIFE recordings could easily be confused with an Aγ-fiber neural response based on latency if using the Erlanger/Gasser scale (Erlanger & Gasser, 1937).

Neuromuscular blockade either eliminated or greatly reduced all EMG responses, as well as the corresponding EMG artifact in the LIFE electrode recordings (n = 7 pigs); aggregate data for all animals can be found in Supplementary Figure 1. It is important to note that although measurements of Aγ, Aδ and B fiber responses in the intact recordings were contaminated by the EMG artifact, the Aβ response in both the intact and neuromuscular junction blockade conditions was unchanged. These data suggest that the dose response curve for Aβ activation, taken via LIFE electrode recordings, were uncontaminated by EMG artifact in the intact state.

### Transection of Somatic Vagus Nerve Branches and the Vagus Trunk to Verify Origins of ENG and EMG

After dose response curves were obtained in both the intact state and after neuromuscular junction blockade, a subset of stimulation amplitudes were repeated in the intact state (post-vecuronium drip) to establish baseline EMG responses immediately before transection of the RL and SL branches was performed. These transections were performed to confirm the RL and SL branches were required for activation of the cricothyroid and cricoarytenoid muscles by VNS. Afterwards, the vagus trunk between the bipolar electrode contacts and LIFEs was also transected to determine whether the recorded ENG signals were indeed compound action potentials evoked by VNS, as opposed to additional unanticipated sources of artifact.

Example ENG and EMG traces before and after the somatic branch transections are shown in Figure 4b. In animals that exhibited a long-latency EMG response and underwent transection of the recurrent laryngeal branch (RLT)(n=9), RLT eliminated the long-latency EMG response in every case (see Supplementary Figure 2). This demonstrates that activation of the cricoarytenoid muscle at low amplitudes was caused by activation of the motor fibers within the stimulation electrode cuff that eventually bifurcate into the recurrent laryngeal branch and innervate the cricoarytenoid muscle. Similarly, in animals that exhibited a short-latency EMG response and underwent transection of the superior laryngeal branch (SLT)(n=8), SLT eliminated the short-latency EMG response in every case (see Supplementary Figure 2). This demonstrates that activation of the cricoarytenoid and cricothyroid muscles at higher amplitudes was caused by activation of the motor fibers outside the stimulation electrode cuff within the nearby superior laryngeal branch which innervates both the cricoarytenoid and cricothyroid muscles. These experimental results confirm that VNS-evoked neck muscle contractions are not due to immediate, direct electrical activation but instead are elicited by VNS-evoked action potentials in vagus nerve fibers that travel along two different length branches resulting in two distinct EMG signals.

Similar to our earlier comparisons between ENG and EMG for neuromuscular junction blockade experiments, elimination of specific signals in EMGs via branch transection also eliminated signals with identical post-stimulus latencies in ENGs; therefore, these transections confirmed that the ENGs were contaminated by EMG artifacts. In animals where the main vagus trunk was transected between the stimulation electrode and the LIFEs (Vagotomy, n=9), transection of the main vagus trunk eliminated all ENG signals. This confirms the remaining LIFE electrode recorded signals prior to vagotomy were indeed neural in origin and not some unanticipated additional artifact (Supplementary Figure 2). The stimulation artifact remained after all transections.

### Stimulation Current Response Curves to Identify ENG and EMG Thresholds for Each Fiber Type

To determine the stimulation threshold for each ENG and EMG response, stimulation-triggered median traces were stacked across stimulation amplitudes to visualize changes in evoked responses (Figure 5a and 5b). For the example in Figure 5, automatic calculation of the rectified area under the curve for each ENG and EMG component within each latency range (i.e., fiber type or EMG component) was used to construct dose-response curves for Aβ- and Aδ/B-fibers (Figure 5c and 5d), as well as long-component and short-component EMGs (Figure 5e and 5f). Visual inspection of Figure 5a suggests thresholds of ∼300, 750, and 2000 µA for activation of Aβ-, Aγ-, and Aδ/B-fibers, respectively, for this example animal. Given temporal overlap of Aβ signals with the larger stimulation artifact, and overlap of Aγ signals with the larger Aβ signals, automatic calculation of amplitude and subsequent determination of threshold was challenging in multiple animals. We therefore used visual identification to produce the data found in Figure 6 and Supplementary Table 1 after generating plots similar to Figure 5a and 5b for ENGs and EMGs in all animals (Supplementary Figure 3).

### Comparison of Thresholds for EMG vs. Activation of Nerve Fiber Types

The first observable ENG component had a latency consistent with Aβ-fibers and a threshold of 300 ± 122 µA. This is substantially lower than the threshold for ENGs with conduction velocities most consistent with parasympathetic efferents (B-fibers) and/or mechanoreceptor afferents (Aδ-fibers) which was 1677 ± 289 µA. Aβ-fiber activation was observed in all animals, while Aδ/B-responses were only observed in 3 of 13 animals despite the use of stimulation amplitudes over twice the average clinically tolerable amplitudes (up to 3 mA vs 1.2-1.3 mA). Consistent with our assumption that the threshold measurement of Aβ-fibers was an indication of Aα-fiber activation at a lower threshold, the threshold for the long-latency EMG (295 ± 180 µA) was slightly lower than the Aβ threshold. The short-latency EMG previously demonstrated to be caused by the activation of the SL branch had a higher threshold (1472 uA ± 643 uA). A higher threshold for SL fibers was expected as the SL is located outside the epineural cuff and activated by current leakage outside the cuff. The thresholds for activation of Aβ-fibers and the long-latency EMG response were generally lower than those thresholds for responses of fiber types associated with desired therapeutic effects, i.e., activation of Aδ/B-fibers (Figure 6).

### Comparison of Thresholds for Stimulation Induced Decreases in Heart Rate

Similar to the frequency of observed Aδ/B fiber responses across the cohort, a bradycardic effect was observed in only six of twelve animals. Notably, the subset of animals in which a bradycardic effect was observed only partially overlapped with the animals in which an Aδ/B fiber response was also detected. The threshold for the bradycardia effect was 1583 ± 1021 µA, which was similar to the average threshold for Aδ/B-fibers across the cohort (See Supplementary Table 1). Interestingly, the bradycardia effect had a lower threshold than the Aδ/B-threshold in some animals, while in others a higher threshold (Supplementary Table 1). Across the 6 animals in which a bradycardic response to VNS was noted, the average threshold for induced bradycardia (1.6 mA) was higher than reported tolerable clinical averages (1.2-1.3 mA) (De Ferrari et al., 2017; Adrian Handforth et al., 1998)

### Comparisons of Vagus Side, Animal Sex, and Stimulus Polarity

Additional analyses were performed to compare ENG and EMG thresholds based on vagus side, animal sex, and stimulus polarity, as well as EMG latencies. Stimulation thresholds for every response except HR were similar between left and right VNS (Figure 6a). However, absolute differences in HR should be interpreted with caution given that isoflurane concentrations were not controlled across animals (see Discussion). No Aδ/B-fiber responses were observed in left side VNS experiments (Figure 6a). Unsurprisingly, the long-latency EMG signal had a shorter latency for right side VNS (Figure 6b) given the right recurrent laryngeal branches more cranially – and is thus a shorter path for motor efferents to travel – than the left recurrent laryngeal (Settell et al., 2019). Thresholds and latencies for every response except HR were similar between male and female pigs (Supplemental Figure 5). Stimulation thresholds for Aβ and EMG responses were almost identical between cathode cranial and cathode caudal configurations (Figure 6c); Aγ and Aδ/B responses were not compared since responses following neuromuscular blockade and following nerve branch transections were not collected for most cathode caudal datasets, and thus may contain motor-evoked potential artifacts at the same latencies as Aγ and Aδ/B responses.

## Discussion

### Placing Results in Pig with Clinical Lead in Context of Historic Large Animal and Clinical Studies

Activation of vagal somatic fibers within or near the vagal trunk has long been speculated to be the source of off-target activation of the muscles of the throat/larynx associated with therapy-limiting side effects (Boon et al., 2009; Castoro et al., 2011). Several methods have been proposed to minimize A-fiber motor activation and maximize Aδ-, B-fiber or C-fiber activation within cervical vagus trunk, including multi-contact electrode arrays to selectively activate certain portions of the vagal trunk (Aristovich et al., 2019) and stimulation waveforms to alter the recruitment order, such as anodal block (Accornero et al., 1977), depolarizing prepulses, slowly rising pulses, quasi-trapezoidal pulses (Fang & Mortimer, 1991; M Tosato et al., 2007), and combinations thereof (Vuckovic et al., 2008). Finite element modeling studies also suggest that unintended collateral activation of the nearby superior laryngeal nerve, exterior to the vagus nerve at the stimulation site, may contribute to off-target effects (Arle et al., 2016). However, prior to the current data, the mechanism of off-target muscle activation had not been carefully verified in a large animal model using an unmodified LivaNova clinical lead.

Somatic fibers within the cuff that eventually branch off into the recurrent laryngeal branch were consistently activated across animals at low amplitudes (0.300 ± 0.122 mA). The threshold for the long latency EMG component via the recurrent laryngeal branch was similar (0.295 ± 0.180 mA). Notably, these stimulation levels are much lower than the average tolerable stimulation amplitudes used clinically (1.2-1.3 mA given similar pulse width) (De Ferrari et al., 2017; A. Handforth et al., 1998). These data suggest that while the activation of the recurrent laryngeal fibers leads to contraction of neck muscles, these contractions do not manifest as the intolerable side effects observed in patients undergoing VNS. However, as human subjects are often ‘ramped up’ slowly over the course of several weeks following initial programming to get to higher tolerable current settings, it is unclear how habituation over time would impact these evoked EMG measures.

In contrast, activation of the somatic fibers within the nearby superior laryngeal branch of the vagus due to current leakage from the cuff is more consistent with the tolerable limits to VNS found in human studies. Initial activation of the cricoarytenoid and cricothyroid muscles through this pathway was often not observed until 0.75 mA and sometimes did not saturate at our maximally applied amplitude of 3 mA, which is more than double tolerable levels in human studies. It should be noted that the LivaNova helical cuff design has relatively little insulation between the edge of each electrode and the edge of the insulating backer (∼1 mm) along the length of the nerve, and consequently may not prevent current leakage as effectively as more extensive epineural cuff designs such as those used in Biocontrol’s Cardiofit human studies (De Ferrari et al., 2017).

The goal of this study was to identify the anatomical sources of off-target muscle activation using the most common FDA-approved clinical electrode in a pig animal model best matching the known diameter of the human cervical vagus, at frequencies commonly used to induce changes in sympathetic/parasympathetic tone. This is in contrast to studies using a similar bipolar spiral cuff design in canines with single stimulation pulses that did not include cardiac responses (Yoo et al., 2013); in comparison to the human cervical vagus, the canine model is notably smaller diameter, has many fewer fascicles, and has a thicker epineurium that increases the distance from the epineural electrodes to the most superficial fibers. As the epineurium is thinner in pigs and humans than in dogs and thus the most superficial fibers are closer to the electrode, one would anticipate that the thresholds for first observable fiber activation to be lower than in canine studies.

Consistent with this hypothesis, the threshold for activation of large diameter A-fibers in the vagal trunks of pigs was smaller (0.3 ± 0.12 mA, 200 µs pulse width) than the threshold found in a canine study (0.37 ± 0.18 mA, 300 µs pulse width) (Yoo et al., 2013). This canine study also used monophasic instead of biphasic stimulation, which should lower the thresholds for activation in comparison to our pig study (Merrill et al., 2005). Similarly, the threshold for the EMG component appearing at the lowest stimulation amplitude in pigs was also found to be smaller (0.3 ± 0.18 mA, 200 µs pulse width) than the threshold in canines (0.36 ± 0.17 mA, 300 µs pulse width). It should also be noted in three of the five dogs tested from Yoo et al. 2013, a notable artifact in the electroneurogram recordings was observed with a short post-stimulus latency signal (3 to 5 ms), similar to the response caused by neck muscle contraction via neurotransmission along the superior laryngeal branch in our pigs. As this response in canines had a short post-stimulus latency and was eliminated following the administration of a neuromuscular blocking agent, the authors hypothesized this component was not related to the activation of the recurrent laryngeal branch. Instead, they proposed this artifact was caused by current spread from the stimulating electrode to a different, unidentified myogenic source (Yoo et al., 2013).

Several important vagus nerve stimulation studies were previously performed in Yorkshire female pigs much larger than those used in our present study (∼100 kg) (M Tosato et al., 2007; Marco Tosato et al., 2006). In these studies, homemade tripolar electrodes were used for stimulation; tripolar stimulation theoretically should increase the rate at which the magnitude of the electric field would decrease with distance from the electrode in comparison to a bipolar configuration, depending on separation between cathode and anodes. However, as the focus of the Struijk studies was to explore closed loop paradigms and novel stimulation pulse strategies using single pulses, data were not collected in such a way that threshold and saturation for A-fiber activation or heart rate responses could be directly compared to our study.

Although the latency range used to define B-fibers differed slightly between the aforementioned studies (M Tosato et al., 2007; Marco Tosato et al., 2006; Yoo et al., 2013), both suggested that the threshold for first observing B-fiber activation was near the border of the average tolerated clinical parameters from human studies of 1.2-1.3 mA given 200 µs pulse widths and 20-30 Hz. Similarly, our results indicate that the threshold for activation of Aδ- or B-fibers was 1.67 ± 0.29 mA (200 µs pulse width). In these prior studies, C-fiber activation was observed only at stimulation parameters well beyond levels that are tolerable by human patients (17 ± 7.6 mA with 300 µs pulse width in Yoo et al. 2013; seen only in two cases by Tosato et al. 2006 at ≥6 mA with 600 µs pulse width). C-fiber activation was not observed in our study^1^, unsurprisingly as 3 mA was the maximal stimulation used. We observed extensive concurrent activation of the neck muscles, which tended to cause electrodes to be displaced from tissue, at amplitudes larger than 3 mA. Considering the previous work exploring higher amplitudes – and the lack of clinical relevance in exploring these values due to generation of intolerable side effects – we concluded that there was limited benefit to exploring higher current amplitudes.

The aggregate data across these studies are difficult to reconcile with epineural cuff recordings performed during human VNS surgery (Evans et al., 2004), which observed evoked compound action potential components with a conduction latency consistent with C-fibers. This putative C-fiber component was observed within 10 ms of stimulation, and in two of the patients the “C-fiber” was apparent at the lowest stimulus setting used (0.2 mA, 200 µs pulse width) (Evans et al., 2004). C-fiber responses were reported in all patients, despite the maximal applied amplitude of 3 mA with a pulse width of 200 to 500 µs. As this study was performed opportunistically during human VNS implantation, understandably neither pharmacological muscle block nor transection of the vagus somatic branches was performed to exclude motor-evoked muscle potentials contaminating the ENG recordings. Data from the canine study in which pharmacological neuromuscular junction blockade was performed (Yoo et al., 2013), and our present study sequentially transecting the recurrent laryngeal and superior laryngeal branches, strongly suggest that this response in the human patients was EMG artifact rather than ENG from C-fibers. We specifically show in our current study that long-latency EMG signal via the recurrent laryngeal pathway can create long-latency artifacts in the ENG recordings.

Collectively, these results provide strong evidence that direct activation of C-fibers is unlikely to be the source of VNS therapeutic effects (Krahl et al., 2001; M Tosato et al., 2007; Marco Tosato et al., 2006; Yoo et al., 2013). Instead, tolerable therapeutic current amplitudes are limited to levels just at the cusp of activating Aδ-fibers (mechanoreceptor afferents) and B-fibers (pre-ganglionic efferents). Thus, tolerable therapeutic current amplitudes are unlikely to be sufficient to activate the entire population of Aδ-and B-fibers within the cervical vagus. These data would help explain the high variability of therapeutic outcomes for VNS in all indications (De Ferrari et al., 2017; A. Handforth et al., 1998; Kimberley et al., 2018; Koopman et al., 2016; Tyler et al., 2017). These data are also consistent with a recent rodent study directly comparing optogenetic and electrical activation of the vagus (Rajendran et al., 2019).

Activation of somatic recurrent laryngeal fibers at low thresholds also has important implications for VNS therapeutic mechanisms. Activation of large diameter efferent fibers and the muscles innervated by those fibers at low levels of stimulation presumably would cause indirect activation of sensory pathways below perceptual thresholds (Bruce & White, 2012) which project to the trigeminal sensory nucleus which also indirectly connects to NTS. This produces a possible confound in VNS studies using indirect surrogate measures of nerve engagement such as functional magnetic resonance imaging (fMRI) or somatosensory evoked potentials (SSEPs). In one study exploring VNS using SSEPs in humans, they noted that VNS SSEPs had four signal peaks and that all but the earliest component disappeared after administration of a muscle relaxant (Usami et al., 2013). Indirect activation of this sensory pathway may also have implications for studies of VNS for plasticity such as the hypothesized vagal pathway to facilitate learning that engages nucleus basalis, locus coeruleus, and other brain areas via the nucleus of the solitary tract (Hays et al., 2013).

### Implications for Non-Invasive Stimulation of the Cervical Vagus

Our results demonstrate that the large diameter efferent fibers within the cervical vagus that eventually become the recurrent laryngeal branch are activated at much lower thresholds than Aδ-, B-, or C-fibers. Additionally, the motor efferent fibers within the nearby superior laryngeal branch were activated near clinically tolerable levels. It is important to note that the superior laryngeal branch of the vagus can form a ramus of communication with the ascending recurrent laryngeal branch of the vagus nerve underneath or near the thyroid cartilage. This is under the location of electrodes for non-invasive VNS, and the distal portions of the superior and recurrent laryngeal branches are much more superficial – closer to the skin of the neck – than the trunk of the cervical vagus nerve.

Activation of the somatic fibers of recurrent and superior laryngeal branches are responsible for activation of the cricoarytenoid and cricothyroid muscles in response to VNS. Therefore, if non-invasive VNS were engaging the pre-ganglionic efferent or sensory afferent fibers of the cervical vagus, presumably one would have to activate these deep muscles of the neck first. However, the off-target effects of invasive VNS including cough, throat pain, voice alteration and dyspnea (De Ferrari et al., 2017; A. Handforth et al., 1998) are not the off-target effects reported for non-invasive VNS. Non-invasive VNS is known to cause lip curl due to activation of the superficial muscles of the neck (Silberstein et al., 2016), which precludes higher levels of stimulation in clinical practice. In short, even the low threshold motor fibers of the deep vagus are not activated by non-invasive VNS. These data suggest that any therapeutic effect of non-invasive VNS is not through direct activation of the fibers at the cervical vagus, but may be achieving its effects through indirect pathways. For example, sinoatrial baroreceptor afferents have been demonstrated to sometimes also travel through the more superficial recurrent laryngeal branch and return to the cervical vagus via the ramus of communication with the superior laryngeal branch (Jacobs et al., 1976; Sanders et al., 1987; Strauss et al., 1973). However, a prior study attempting transcutaneous stimulation of the superficial portion of the recurrent laryngeal branch in monkey demonstrated closing of the glottic aperture due to activation of the cricoarytenoid and cricothyroid muscles, but without any cardiac effects (Sanders et al., 1987).

### Study Limitations

The isoflurane anesthesia used in our experiments should only impact synaptic transmission (Baumgart et al., 2015; Herring et al., 2009) and would not change the thresholds for direct electrical activation of vagus fibers. Likewise, fentanyl analgesia is known to cause bradycardia, which is dependent on vagal-mediated central pathways and may have modulated both the effects of isoflurane and VNS (Laubie et al., 1977; Reitan et al., 1978). Isoflurane is commonly used in vagus and carotid sinus nerve stimulation studies (Georgakopoulos et al., 2009; Thompson et al., 1998), and is not known to impact the stimulation thresholds for motor efferent fibers to elicit muscle contraction. Isoflurane is known to blunt baroreflex mediated cardiac responses (Kotrly et al., 1984; Seagard et al., 1983); however, baroreceptor-induced heart rate changes can still be observed at isoflurane concentrations up to 2.1% (Bagshaw & Cox, 1988). In our study, isoflurane levels were adjusted through the duration of the experiment depending on plane of anesthesia. While all efforts were made to keep the isoflurane concentration as low as possible, isoflurane concentrations at or above 2% were occasionally needed. As cardiac effects were not the primary outcome for these studies, a careful analysis of cardiac response based on isoflourane concentration was not performed. Although robust stimulation-evoked bradycardia was observed at isoflurane concentrations of 2-2.5%, anecdotally these responses were generally smaller than at lower isoflurane concentrations in the same animal. The synaptic blunting caused by isoflurane anesthesia may require activation of a larger number of parasympathetic efferent fibers to the heart or mechanoreceptor afferent fibers to elicit a heart rate response, and therefore change the relationship between the threshold for observing the compound action potential in the neural recordings and the threshold for an associated heart rate response. An additional consideration, due to the sparse sampling nature of the LIFEs for compound action potential measurements, is that in some of our experiments the LIFEs may have been placed directly inside or adjacent to fascicles containing parasympathetic efferent fibers and may thus cause the apparent threshold for those fibers to be lower than an experiment where the LIFEs were placed in or near fascicles without those fibers. Likewise, these considerations likely contributed to our observation that the apparent threshold for Aδ/B-fibers did not correlate to the apparent threshold for the bradycardia effect on an animal by animal basis (Supplementary Table 1).

Moreover, as these experiments were acute, there are several variables that may impact the spatial distribution of applied currents and thus thresholds. The acute surgical trauma often resulted in edema creating fluid at or near the electrode/nerve interface that was tracked visually and removed with surgical gauze when noted. The required cut down and associated surgical pocket displaces tissue in the region nearby the electrode, and it is uncertain if motor nerve fibers other than those of the recurrent laryngeal and superior laryngeal were unintentionally removed from the vicinity of the electrodes. Perhaps most notably, the acute surgical environment does not approximate well the fibrous scar that forms around chronically implanted electrodes, and this would presumably increase the distance of the stimulation electrodes to the nerve trunk.

Finally, the superior laryngeal branch in the pig is located more caudal than human (Settell et al., 2019). In the pig, the superior laryngeal branches within the common surgical window for vagus nerve stimulator implants, whereas in the human the superior laryngeal branches much closer to the mandible. However, the external branch of the superior laryngeal in the human traverses near the carotid bifurcation and may be affected similarly to the superior laryngeal in pig. Despite these potential limitations, the acute experimental results presented here were consistent with finite element model predictions of human vagus nerve activation (Arle et al., 2016) and stimulation parameters reported tolerable prior to side effects putatively associated with neck muscle contraction in humans (De Ferrari et al., 2017).

### Final Thoughts and Next Steps

The FDA-approved human VNS lead by LivaNova consists of two electrodes with active contacts that wrap approximately 270° around the surface of the nerve trunk in a helical cuff, and there is a relatively short distance between the edge of the electrode and the edge of the insulating backer along the length of the nerve. Simple solutions such as increasing the size of the insulating backer or altering the path of the descending external branch of the superior laryngeal during surgery may be sufficient to minimize off-target effects mediated by activation of the superior laryngeal branch in passing.

Avoiding activation of somatic recurrent laryngeal fibers within the cervical vagus nerve in the cuff may be more challenging. One potential solution is a multi-contact array of smaller electrodes (Aristovich et al., 2019) predicated on the idea that there is functional organization of the fibers within the cervical vagus trunk (Settell et al., 2019), termed ‘vagotopy’, that can be leveraged to better isolate vagal fibers associated with specific therapeutic effects. Although there have been demonstrations using multi-contact vagus nerve cuffs in small and large animal models to differentiate between activation of cardiac pathways versus respiratory pathways (Aristovich et al., 2019), it is unknown if there is sufficient functional vagotopy within the cervical vagus nerve to avoid the common therapy-limiting side effect of deep neck muscle activation. Detailed histology of the vagus across animal models and in human subjects has been recently performed (Settell et al., 2019), and additional work to determine exact fiber compositions and locations are warranted, as the specifics may help inform the most simple electrode design in terms of contact size, spacing, overall number and orientation of electrodes that can improve isolation of specific vagal pathways.

The use of tailored stimulation strategies such as anodal block, hyperpolarizing pre-pulses, and guard anodes were proposed to change the ratio of activation between A-, B-, and C-fibers, with some promising results in acute animal models (M Tosato et al., 2007; Marco Tosato et al., 2006; Vuckovic et al., 2008). However, these strategies have not been implemented in clinical settings. This may in part due to the fundamental difficulty in translating new technology into clinical settings. However, these solutions may also be less suitable when scaling to more complex large diameter nerves in the human or large animal model under chronic conditions. In general, as the size of the nerve trunk increases, the distance from the electrode to the fibers of interest increases and spans a larger range, and the thresholds for activation between cathodic or anodic leading biphasic waveforms becomes difficult to distinguish. Consistent with this premise, both in prior canine studies (Yoo et al., 2013) and in the present study, no obvious clinically relevant difference was observed in the thresholds for activation when reversing the location of the cathode and anode.

## Conclusion

Side effects of VNS present a significant barrier to therapeutic outcomes in the clinic. To better understand the source of these side effects, we stimulated the human-sized cervical vagus of domestic pigs using the same bipolar helical lead used clinically. The cricoarytenoid muscle was activated via motor fibers running within the cuff which eventually become part of the recurrent laryngeal branch at very low thresholds (∼0.3 mA). At higher levels of stimulation (∼1.4 mA) approaching clinically tolerable limits, current leakage outside of the cuff activated the motor fibers in the nearby superior laryngeal branch, causing contraction of both the cricoarytenoid and cricothyroid muscles. Stimulation at the average tolerable levels derived from clinical studies (∼1.3 mA) was often insufficient to activate Aδ- and/or B-fibers and/or evoke bradycardia, and Aδ-/B-fiber activation and bradycardia were not observed in multiple animals despite stimulation amplitudes as high as 3 mA. Our data also suggest that previously reported C-fiber recordings were due to artifacts arising from EMGs elicited by activation of short- and long-latency motor pathways through the superior and recurrent laryngeal branches of the vagus, respectively.

Collectively, these data suggest that the superior laryngeal branch of the vagus nerve may be an anatomical landmark that should be avoided during VNS. Moreover, strategies to avoid the therapy-limiting side effect, such as use of high density epineural electrodes to take better advantage of functional organization within the cervical vagus, should be explored. In addition, mechanisms of VNS that do not depend on direct activation of sensory afferents from the visceral organs should be investigated with increased attention.

## Acknowledgements

The authors would like to acknowledge funding from The Defense Advanced Research Projects Agency (DARPA) Biological Technologies Office (BTO) Targeted Neuroplasticity Training Program under the auspices of Doug Weber and Tristan McClure-Begley through the Space and Naval Warfare Systems Command (SPAWAR) Systems Center with (SSC) Pacific grant no. N66001-17-2-4010, NIH SPARC OT2 OD025340, and CTSA Grant Number TL1 TR002380 from the National Center for Advancing Translational Science (NCATS). Its contents are solely the responsibility of the authors and do not necessarily represent the official views of the NIH.

We would like to thank Dr. Jamie J. Van Gompel for guidance on placement of the LivaNova lead in our experiments.

## Conflict of Interest Statement

JW and KAL are scientific board members and have stock interests in NeuroOne Medical Inc., a company developing next generation epilepsy monitoring devices. JW also has an equity interest in NeuroNexus technology Inc., a company that supplies electrophysiology equipment and multichannel probes to the neuroscience research community. KAL is also paid member of the scientific advisory board of Cala Health, Blackfynn, Abbott and Battelle. KAL also is a paid consultant for Galvani and Boston Scientific. KAL is a consultant to and co-founder of Neuronoff Inc.

None of these associations are directly relevant to the work presented in this manuscript.

## Footnotes

1 In one animal, a possible C-fiber signal was observed that persisted following transection of both recurrent and superior laryngeal branches; however, this signal resembled the EMG artifacts we identified throughout the cohort and vecuronium was not used in this animal. Thus we hesitated to report the signal as a C-fiber. Please see Supplementary Figures 2 and 3, animal 190403 to observe this example.

