## Supplemental Table and Figures for "Sources of Off-Target Effects of Vagus Nerve Stimulation Using the Helical Clinical Lead in Domestic Pigs"

#### Supplementary Table 1

| | 181205 | 181211 | 190109 | 190122 | 190123 | 190129 | 190227 | 190403 | 190410 | 190501 | 190522 | 190529 |
|---|---|---|---|---|---|---|---|---|---|---|---|---|
| **Vagus Nerve Side** | Left | Left | Right | Right | Right | Right | Left | Left | Left | Right | Right | Right |
| **Animal Sex** | M | M | M | M | M | F | F | F | M | M | F | F |
| **Aβ Threshold (uA)** | 300 | 200 | 300 | 300 | 100 | 200 | 200 | 500 | 250 | 350 | 500 | 400 |
| **Aγ Threshold (uA)** | 1000 | 1000 | 1500 | 750 | 200 | 400 | 300 | 1500 | 450 | - | - | - |
| **Aδ/B Threshold (uA)** | - | - | - | 2000 | 1500 | 1500 | - | - | - | - | - | - |
| **Short EMG Threshold (uA)** | 2000 | - | - | 750 | 2000 | - | 1000 | 1500 | 2500 | 1500 | 500 | 1500 |
| **Long EMG Threshold (uA)** | - | 200 | 200 | 200 | 150 | 200 | 100 | 750 | 350 | 350 | 400 | 350 |
| **HR Threshold (uA)** | 500 | 500 | - | 1500 | - | 3000 | 2500 | 1500 | - | - | - | - |
| **Aβ Velocity (m/s)** | 42.32 | 53.65 | 50.80 | 51.32 | 51.01 | 52.27 | 43.35 | 50.47 | 45.94 | 50.27 | 51.33 | 54.77 |
| **Aγ Velocity (m/s)** | 26.30 | 36.91 | - | 29.81 | 37.12 | 30.09 | - | 39.64 | 31.92 | - | - | 39.67 |
| **Aδ/B Velocity (m/s)** | - | - | - | 11.67 | 17.94 | 14.67 | - | - | - | - | - | - |
| **Short EMG Latency (ms)** | - | - | - | 3.7 | 4.8 | - | 4.4 | 4.8 | 6.4 | 4.8 | 5.6 | 4.6 |
| **Long EMG Latency (ms)** | - | 9.8 | 7.1 | 6.6 | 7.0 | 6.2 | 9.3 | 10.9 | 12.2 | 8.0 | 8.1 | 7.1 |
| **Cranial E to SL Insert (cm)** | 0.0 | 0.2 | 0.4 | 0.5 | 0.4 | 0.4 | 0.3 | 0.8 | 0.6 | 0.8 | 0.5 | 0.7 |
| **Caudal E to SL Insert (cm)** | 1.0 | 1.0 | 1.1 | 0.9 | 1.0 | 0.9 | 1.1 | 1.3 | 1.3 | 1.5 | 1.0 | 1.5 |
| **Cranial E to Center LIFE (cm)** | 6.5 | 6.3 | 8.6 | 9.3 | 9.4 | 8.4 | 9.5 | 9.2 | 10.1 | 8.7 | 8.2 | 8.5 |
| **Caudal E to Center LIFE (cm)** | 5.5 | 5.5 | 7.9 | 8.9 | 8.8 | 7.9 | 8.7 | 8.7 | 9.4 | 8.0 | 7.7 | 7.7 |
| **Cranial E Recurrent Path Length (cm)** | - | - | - | - | - | - | 28.9 | 36.2 | 38.1 | 22.6 | 21.3 | 23.1 |
| **Caudal E Recurrent Path Length (cm)** | - | - | - | - | - | - | 28.1 | 35.7 | 37.4 | 21.9 | 20.8 | 22.3 |
| **Superior Laryngeal Length (cm)** | - | - | - | - | - | - | 3.0 | 3.0 | 4.0 | 2.5 | 3.0 | 2.3 |

| | Avg | Std | M Avg | M Std | F Avg | F Std | L Avg | L Std | R Avg | R Std |
|---|---|---|---|---|---|---|---|---|---|---|
| **Vagus Nerve Side** | - | - | - | - | - | - | - | - | - | - |
| **Animal Sex** | - | - | - | - | - | - | - | - | - | - |
| **Aβ Threshold (uA)** | 300 | 122 | 257 | 84 | 360 | 152 | 290 | 124 | 307 | 130 |
| **Aγ Threshold (uA)** | 789 | 494 | 817 | 459 | 733 | 666 | 850 | 482 | 713 | 572 |
| **Aδ/B Threshold (uA)** | 1667 | 289 | 1750 | 354 | 1500 | N/A | - | - | 1667 | 289 |
| **Short EMG Threshold (uA)** | 1472 | 643 | 1750 | 661 | 1125 | 479 | 1750 | 645 | 1250 | 612 |
| **Long EMG Threshold (uA)** | 295 | 180 | 242 | 86 | 360 | 248 | 350 | 286 | 264 | 99 |
| **HR Threshold (uA)** | 1583 | 1021 | 833 | 577 | 2333 | 764 | 1250 | 957 | 2250 | 1061 |
| **Aβ Velocity (m/s)** | 49.79 | 3.88 | 49.33 | 3.86 | 50.44 | 4.28 | 47.15 | 4.81 | 51.68 | 1.49 |
| **Aγ Velocity (m/s)** | 33.93 | 5.05 | 32.41 | 4.66 | 36.47 | 5.52 | 33.69 | 5.87 | 34.17 | 4.99 |
| **Aδ/B Velocity (m/s)** | 14.76 | 3.14 | 14.80 | 4.43 | 14.67 | N/A | - | - | 14.76 | 3.14 |
| **Short EMG Latency (ms)** | 4.9 | 0.8 | 4.9 | 1.1 | 4.9 | 0.5 | 5.2 | 1.0 | 4.7 | 0.7 |
| **Long EMG Latency (ms)** | 8.4 | 1.9 | 8.5 | 2.2 | 8.3 | 1.8 | 10.5 | 1.3 | 7.2 | 0.7 |
| **Cranial E to SL Insert (cm)** | 0.5 | 0.2 | 0.4 | 0.3 | 0.5 | 0.2 | 0.4 | 0.3 | 0.5 | 0.2 |
| **Caudal E to SL Insert (cm)** | 1.1 | 0.2 | 1.1 | 0.2 | 1.2 | 0.2 | 1.1 | 0.1 | 1.1 | 0.3 |
| **Cranial E to Center LIFE (cm)** | 8.5 | 1.1 | 8.4 | 1.4 | 8.7 | 0.6 | 8.3 | 1.8 | 8.7 | 0.4 |
| **Caudal E to Center LIFE (cm)** | 7.9 | 1.2 | 7.7 | 1.6 | 8.1 | 0.5 | 7.6 | 1.9 | 8.1 | 0.5 |
| **Cranial E Recurrent Path Length (cm)** | 28.4 | 7.3 | 30.3 | 10.9 | 27.4 | 6.7 | 34.4 | 4.8 | 22.3 | 0.9 |
| **Caudal E Recurrent Path Length (cm)** | 27.7 | 7.3 | 29.7 | 11.0 | 26.7 | 6.8 | 33.7 | 5.0 | 21.7 | 0.8 |
| **Superior Laryngeal Length (cm)** | 3.0 | 0.6 | 3.3 | 1.1 | 2.8 | 0.3 | 3.3 | 0.6 | 2.6 | 0.4 |

Supplementary Table 1: Key anatomical distances, electrode locations, and ENG and EMG summaries. Entries that contain a dash ("-") denote the following: for thresholds, ENG or EMG was not observed or the data was unclear; for velocities and latencies, threshold could not be determined and these cannot be calculated; for distances, data was never collected. Definitions: **Short-latency EMG**, (See Figure 2B). **Long-latency EMG**, (See Figure 2B). **Cranial (or caudal) E to SL Insert**, distance from each stimulation electrode contact (more cranial or more caudal electrode contact) to the point where the superior laryngeal (SL) branch leaves the main vagus trunk. **Cranial (or caudal) E to Center LIFE**, distance from each stimulation electrode contact to the center of the LIFE cluster. All ENG summaries for a given animal are the mean average of the response type from all LIFEs that have observable signal. **Cranial (or caudal) E Recurrent Path Length**, distance from each stimulation electrode contact to the cricoarytenoid muscle along the path of the main vagus trunk and recurrent laryngeal branch. **Superior Laryngeal Length**, distance from the point at which the superior laryngeal branch leaves the main vagus trunk to where the superior laryngeal branch innervates the cricothyroid muscle. Average (Avg), Standard Deviation (Std), Female (F), Male (M), Left Vagus (L), Right Vagus (R).

**Supplementary Figure 1**

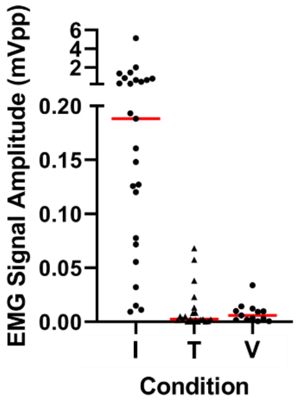


Supplementary Figure 1: Summary of EMG signal amplitudes, both early and late components combined, before and after transections, as well as vecuronium. Application of vecuronium (V) or transection of the recurrent laryngeal and superior laryngeal branches (T) eliminates most EMG signals.

**Supplementary Figure 2**

ENG and EMG traces for branch and trunk transections in all animals. See diagram a for abbreviations. See diagram b for explanation of panel organization for each animal.

Transection of the two somatic branches of the vagus were performed to test the hypothesis that vagus nerve stimulation causes action potentials that travel along the branches of the vagus into neck muscles, thus causing the apparent VNS-evoked neck muscle contractions. Transection of the recurrent laryngeal **(RLT)** always removed the long latency EMG component and ENG signals with identical latencies (EMG artifacts). Transection of the superior laryngeal **(SLT)** always removed the short latency EMG component and ENG signals with identical latencies (EMG artifacts). These experiments confirmed that VNS-evoked neck muscle contractions (and corresponding EMG signals) occur due to action potential signaling along the vagus nerve branches, as opposed to direct muscle activation. These experiments also suggest that neck muscle contractions can create EMG artifacts in ENGs that might be mistaken for nerve fiber signals.

Transection of the main vagus trunk after transection of the vagus somatic branches was performed to confirm that remaining ENG signals were caused by action potentials elicited at the stimulation electrode cuff that move down the vagus to where the LIFEs were located, as opposed to some unidentified artifact. Transection of the vagus trunk cranial **(CrT)** to the stimulation electrode had no effect on any signals recorded. This makes sense because action potentials generated under the cuff could still travel caudal to the cuff, which is where the LIFEs were located. Transection of the vagus trunk caudal **(CaT)** to the stimulation electrode always removed all remaining signals in the ENG recordings. This suggests the ENG signals recorded at the LIFEs prior to transection of the main vagus trunk were indeed action potentials evoked by electrical stimulation at the stimulation electrode cuff.

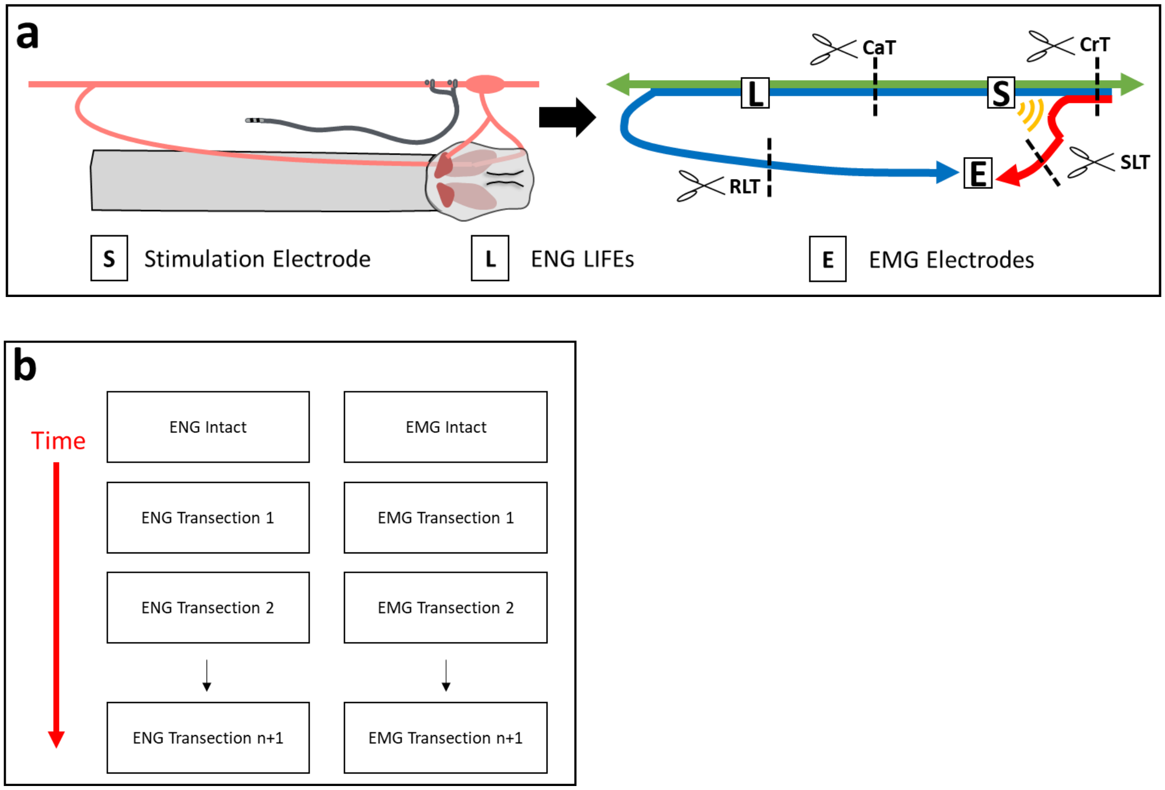


Supplementary Figure 2 Diagrams: **a)** Left) cartoon of surgical field, right) wiring diagram of fibers (colored arrows) with electrode locations (lettered boxes) and transection locations (scissors). Blue arrow is fibers of the recurrent laryngeal branch, red arrow is fibers of the superior laryngeal branch, green arrow is all other fibers. Yellow semi-circles is the expected current leakage from the stimulating electrode to the superior laryngeal branch outside the cuff. Black dashed lines with scissors are the locations of branch or nerve transections, with corresponding abbreviations. Recurrent laryngeal branch transection (RLT), superior laryngeal branch transection (SLT), cranial to stimulation electrode vagus trunk transection (CrT), caudal to stimulation electrode vagus trunk transection (CaT). **b)** Organization of example data for each animal. Left panels are ENG, right panels are simultaneously collected EMG. Colored lines in ENG data are different electrodes, colored lines in EMG data are collected from the cricothyroid muscle (blue lines) and the cricoarytenoid muscle (red lines). The top two panels are collected with no transections (intact). Moving from top to bottom, the panels are collected in order through time. Some electrode traces were removed in some animals due to excessive noise, artifacts, or drift.

**181205**

RLT not performed.

Only animal with no apparent long component EMG. Since we did not isolate the recurrent laryngeal branch for transection, there is a chance the recurrent was severed accidentally during surgical cutdown.

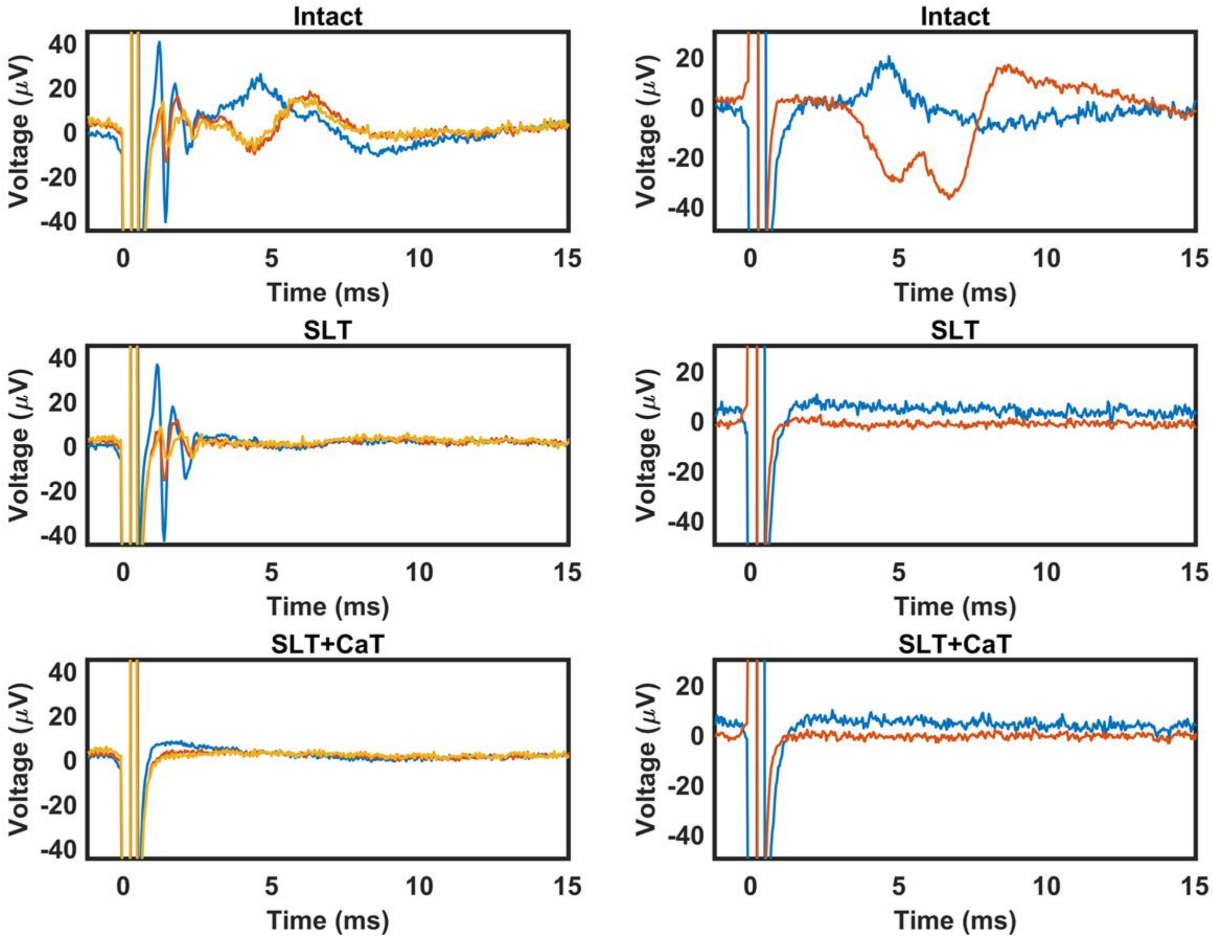

**181211**

No transections were performed in this animal.

**190109**

No short component EMG in this animal.

Clear long component EMG artifact in the ENG signals.

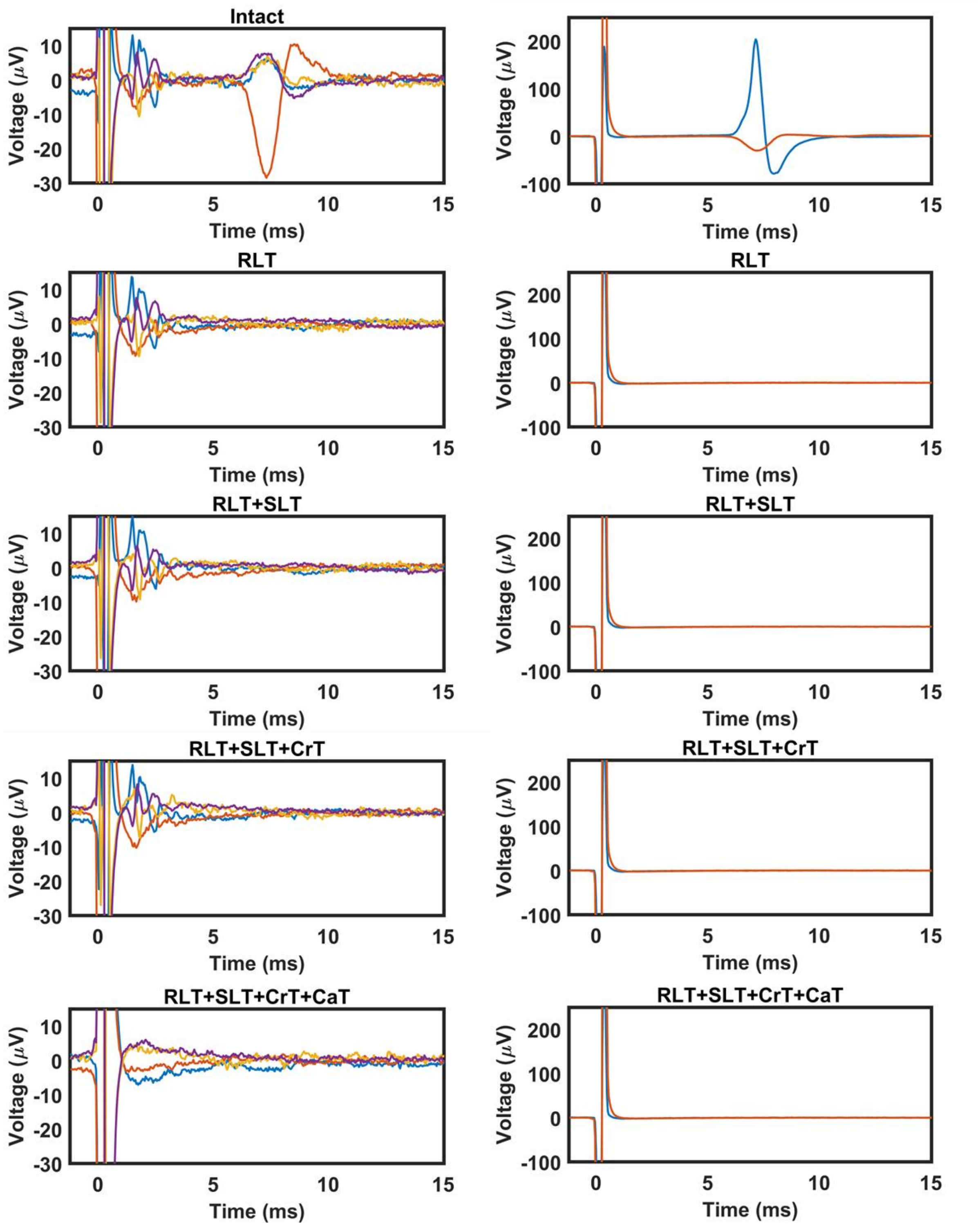

**190122**

Long component EMG is small but present.

Good example of how the long latency EMG causes an artifact in the same temporal location of Aδ/B- signals; compare “Intact” to “RLT+SLT”.

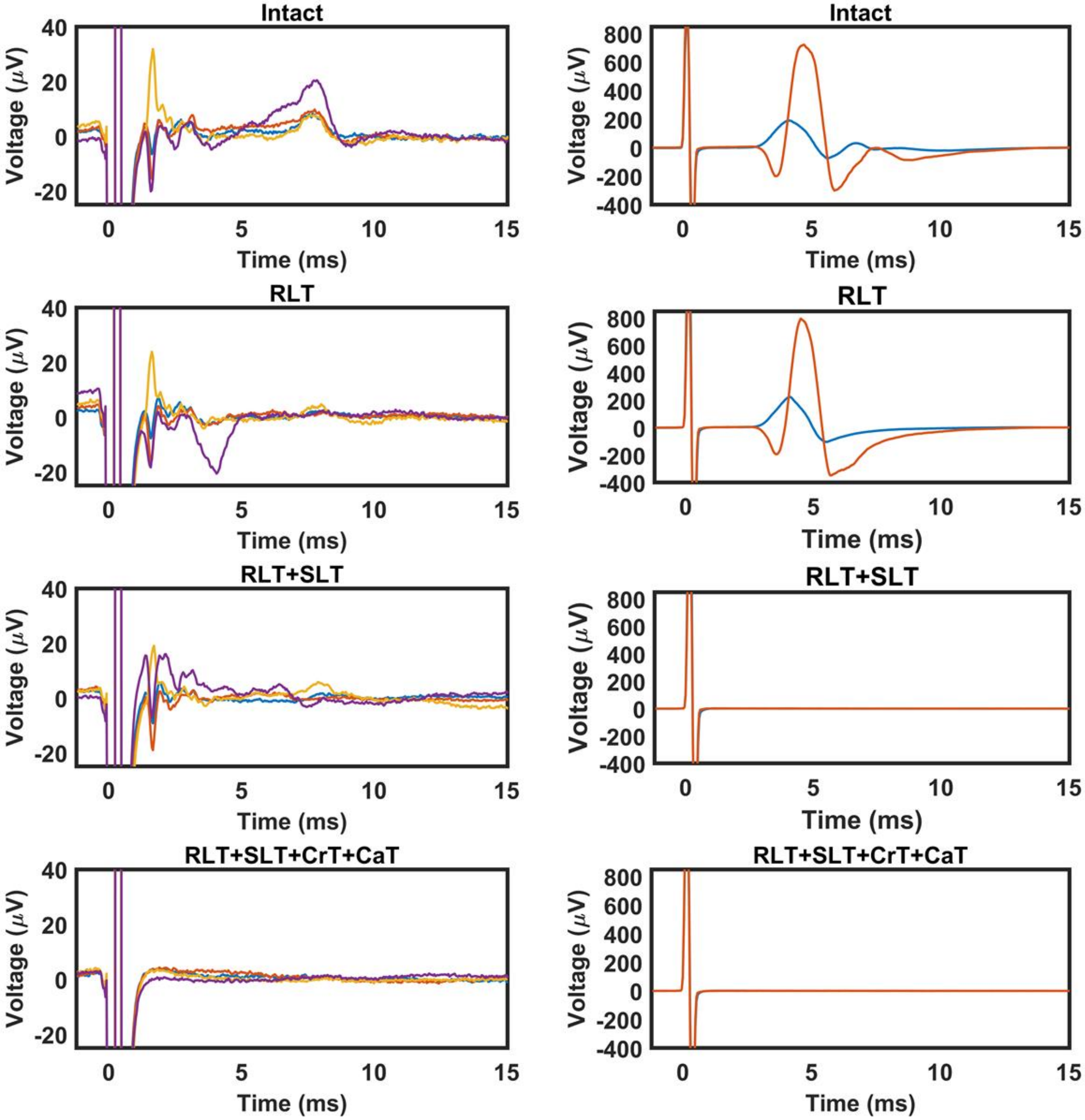

**190123**

Short component EMG is small but present, see zoom below comparing EMG “RLT” to “RLT+SLT”.

Good example of how sometimes no EMG artifacts will happen in ENG signals despite large EMG components.

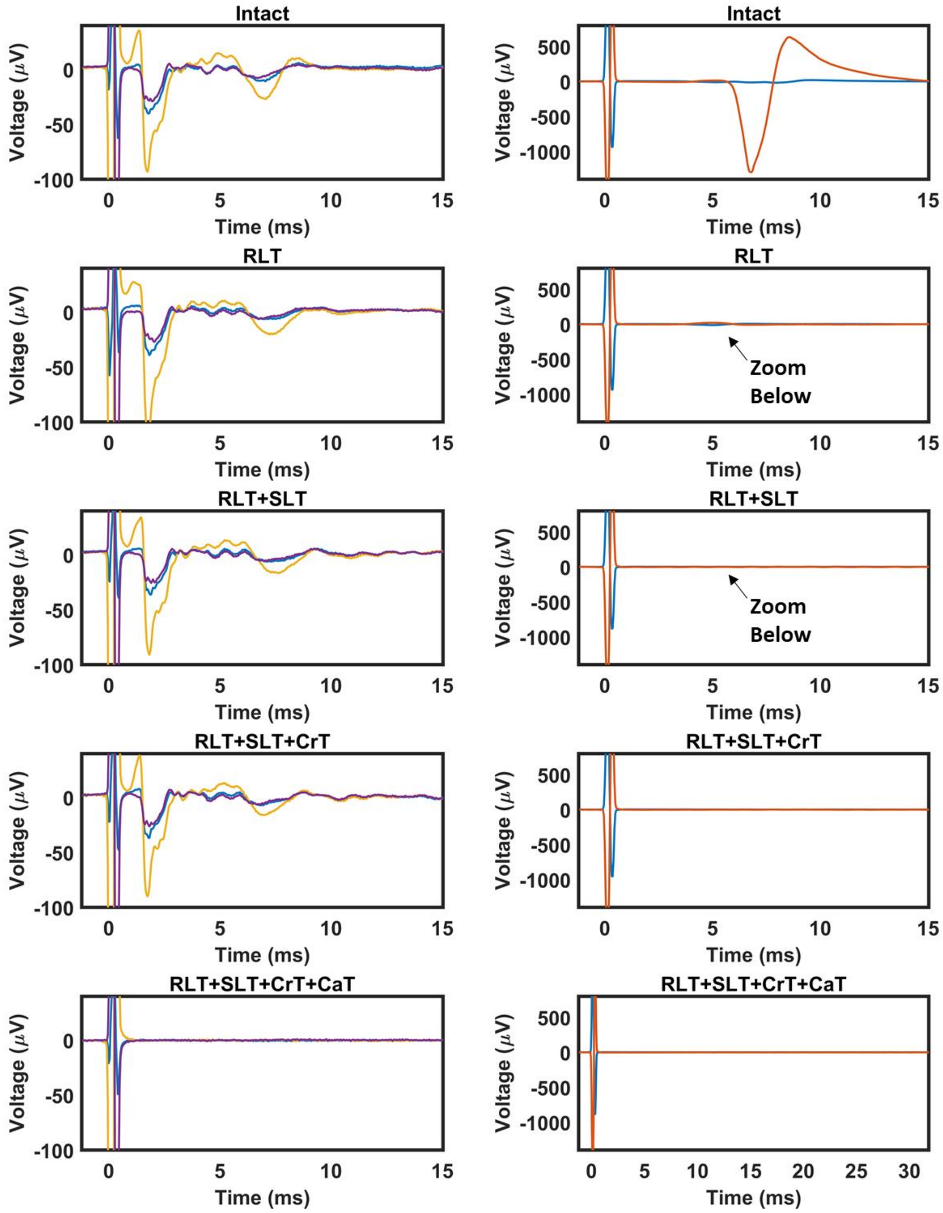

**190123 EMG Amplitude Zooms**

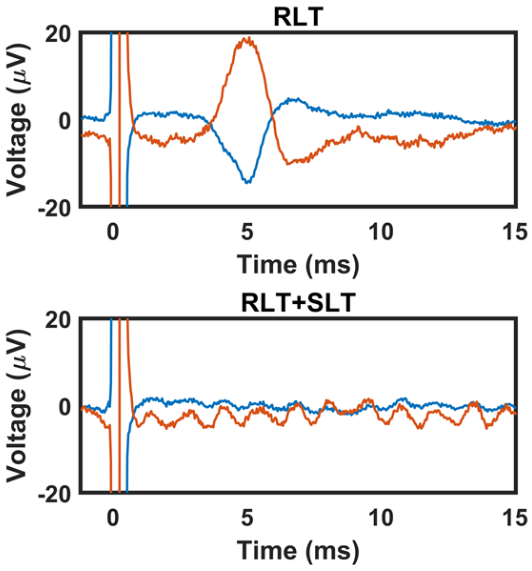

**190129**

No short component EMG in this animal.

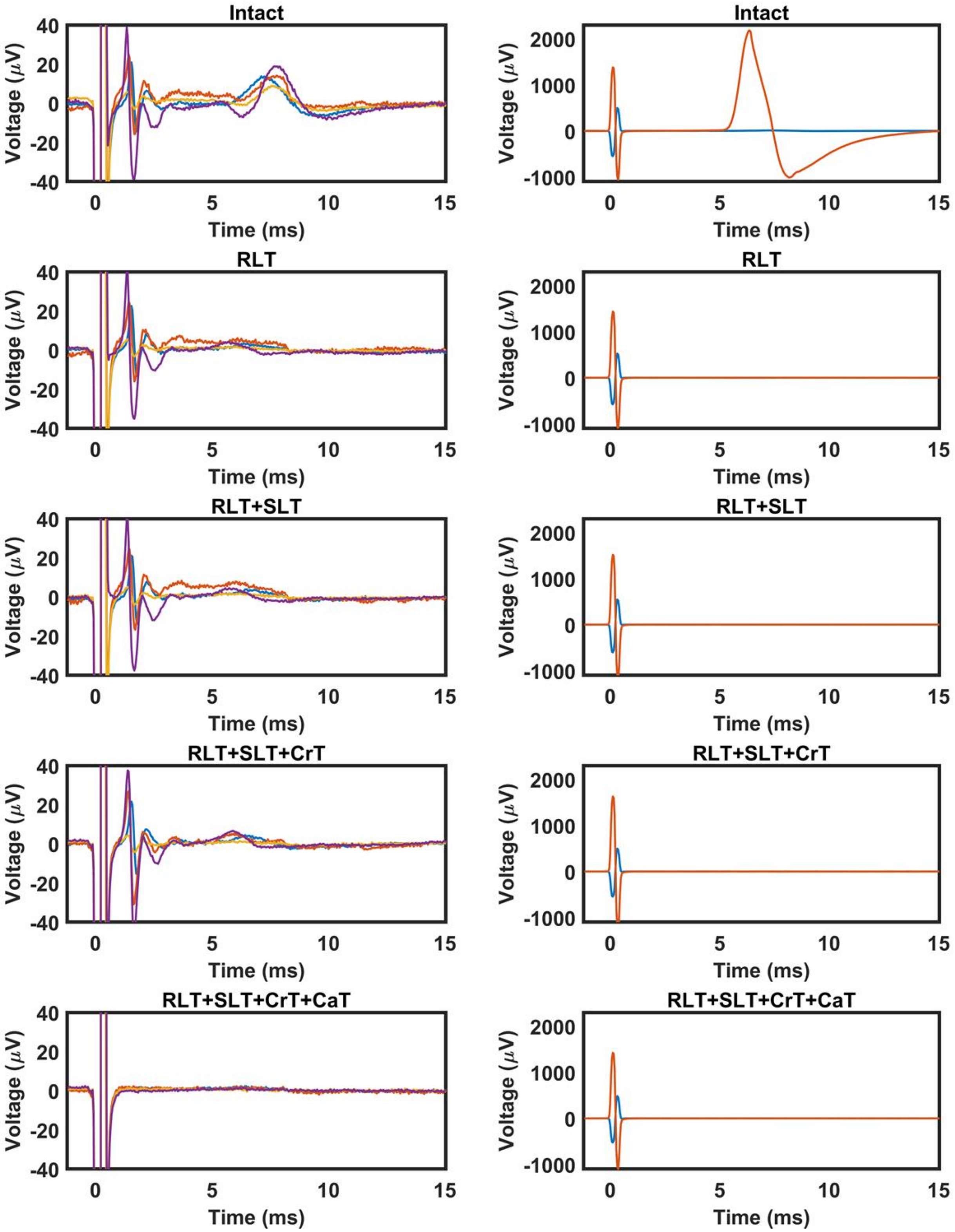

190227

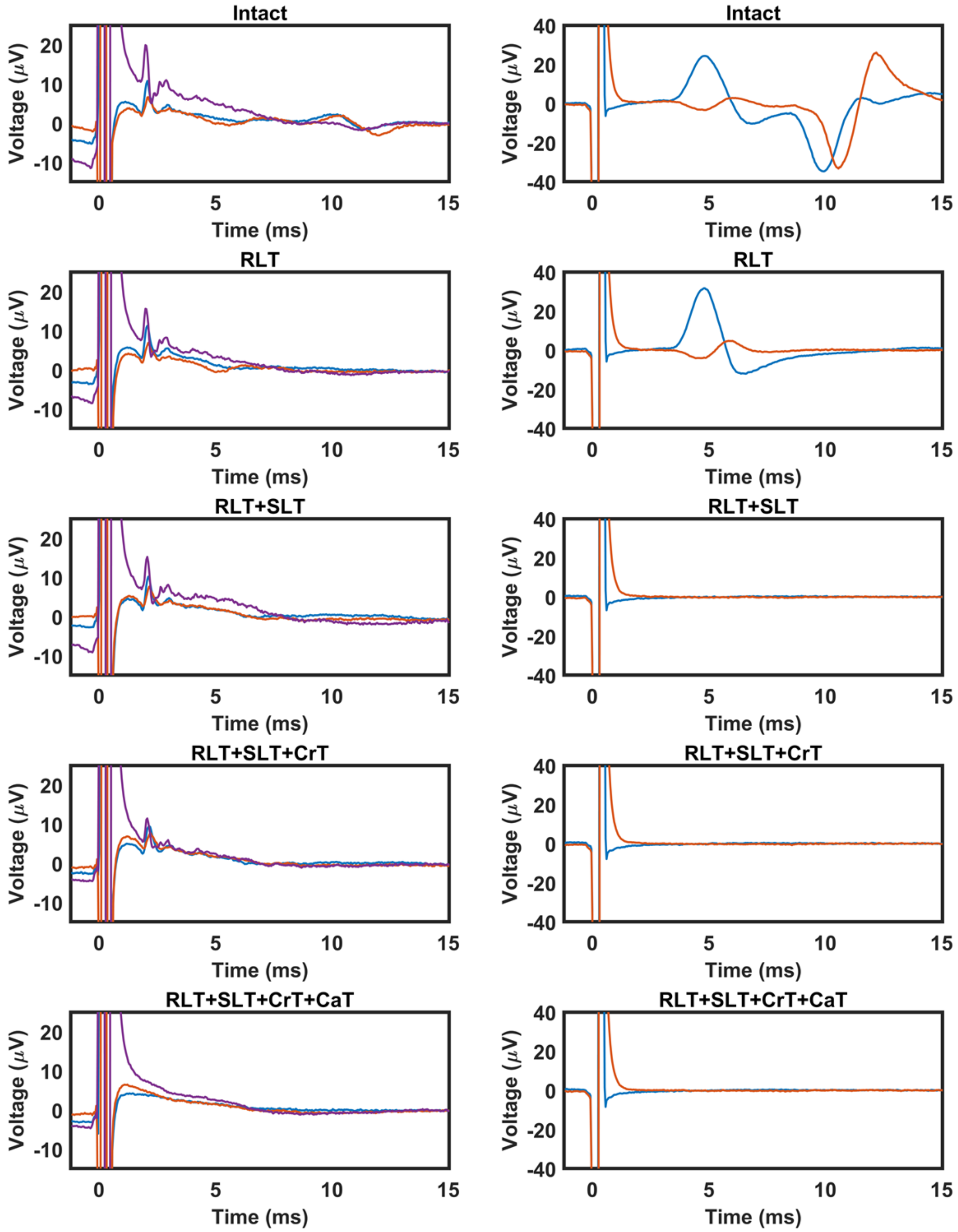

Intact
RLT
RLT+SLT
RLT+SLT+CrT
RLT+SLT+CrT+CaT
Voltage (μV)
Time (ms)

**190403**

Two temporal zooms are presented. First, 0 to 15 seconds to match the rest of the animals. Second, 0 to 30 seconds to show the possible C-fiber, which was not observed in any of the other animals. Note that the possible C-fiber remains after both branch transections and is eliminated following transection of the main vagus trunk caudal to the stimulating electrode.

Only animal that SLT was performed before RLT. Respective effects on short and long component EMG signals is the same as RLT then SLT. Long component EMG is small but present; see zoom comparing “SLT” to “SLT+RLT” below.

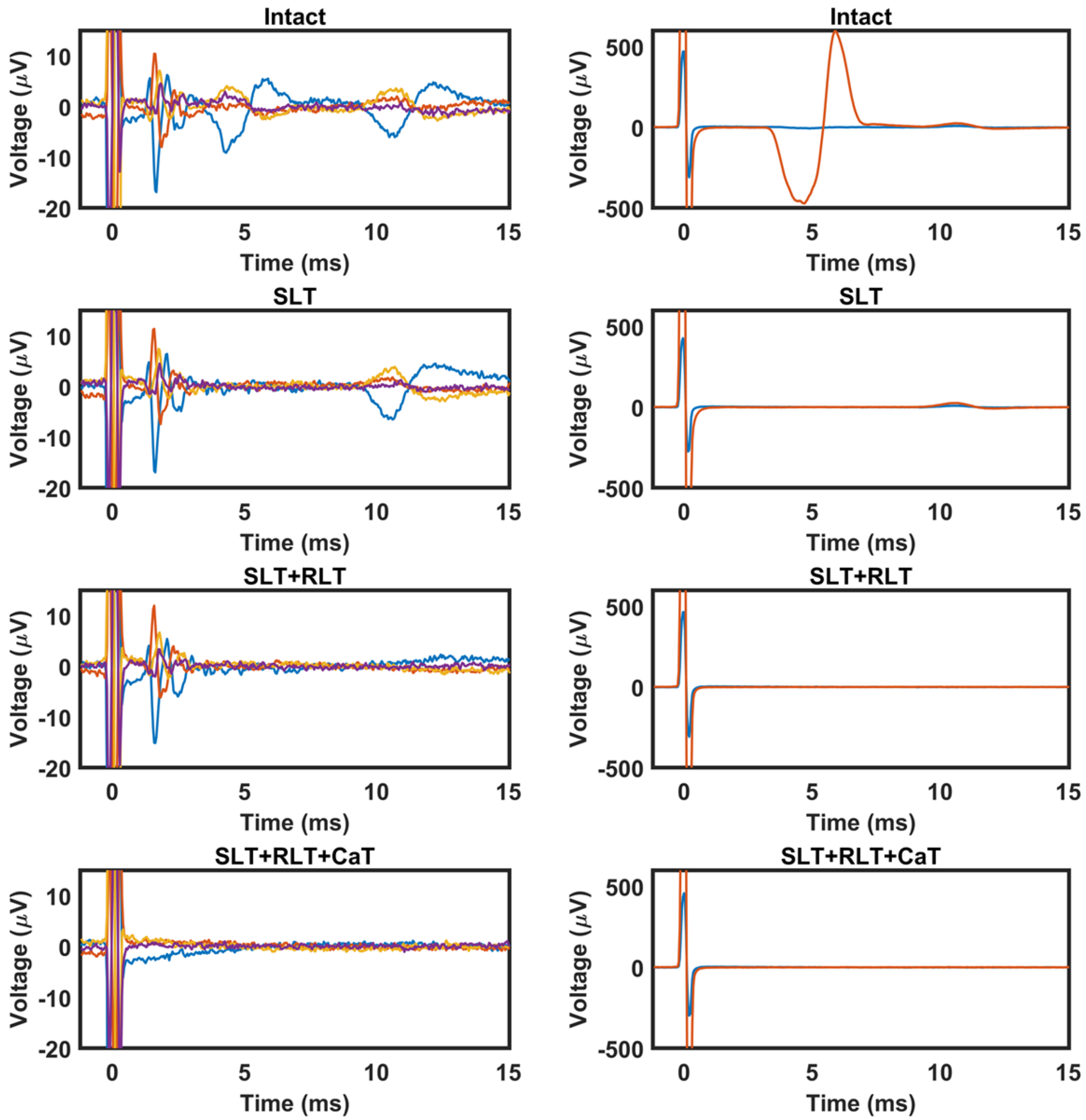

**190403 EMG Amplitude Zooms**

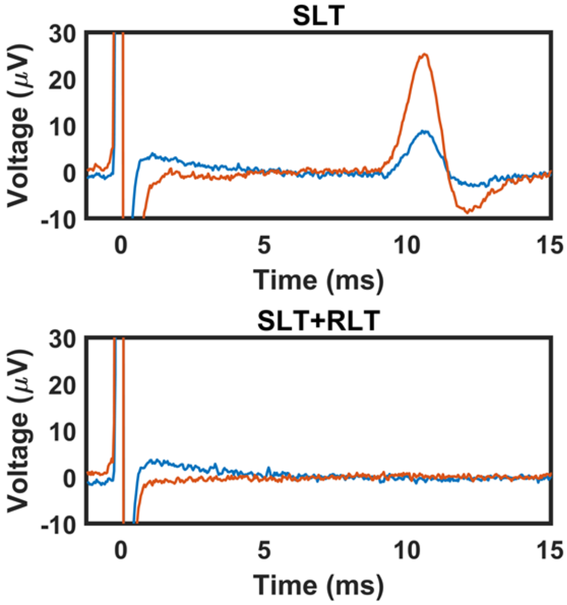

**190403 Possible C-fiber**

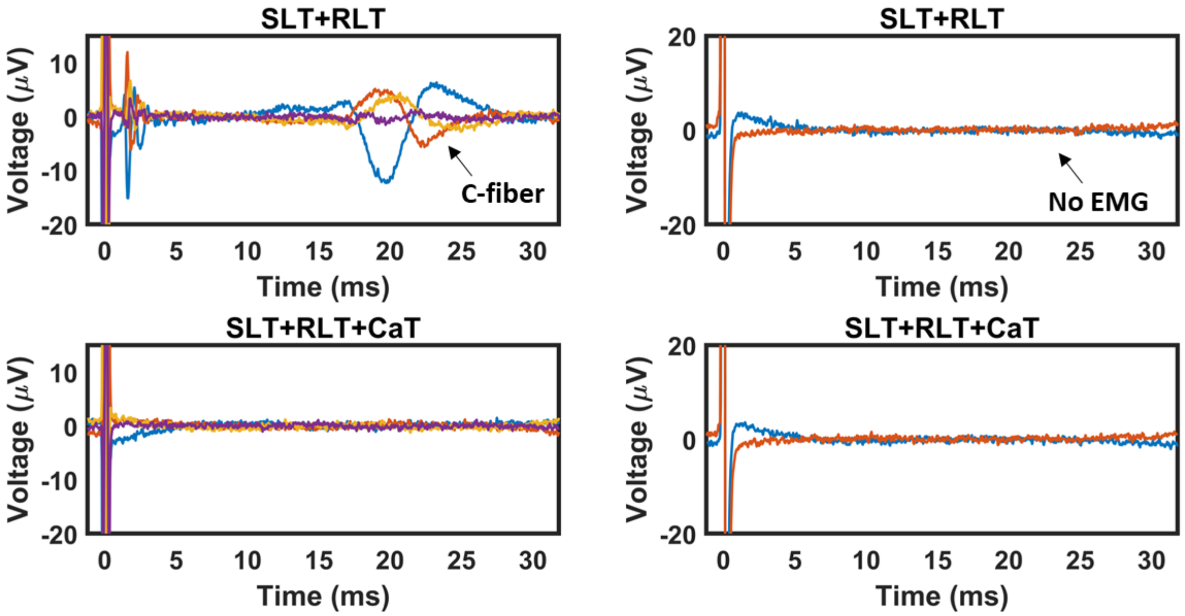

**190410**

Main vagus trunk transection not performed.

Branch transections not performed.

**190501**

Main vagus trunk transection not performed.

Vecuronium was still in effect during branch transection data.

**190522**

ENG fiber signals were not clear in the intact condition for this animal prior to transections.

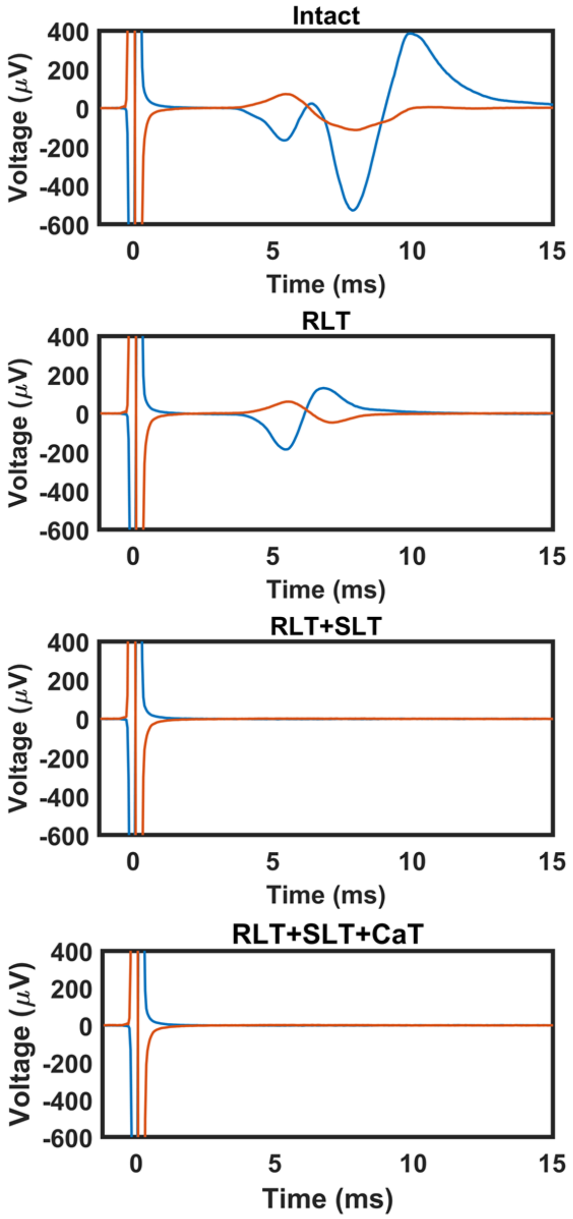

**190529**

Subtracted the second channel from the others.

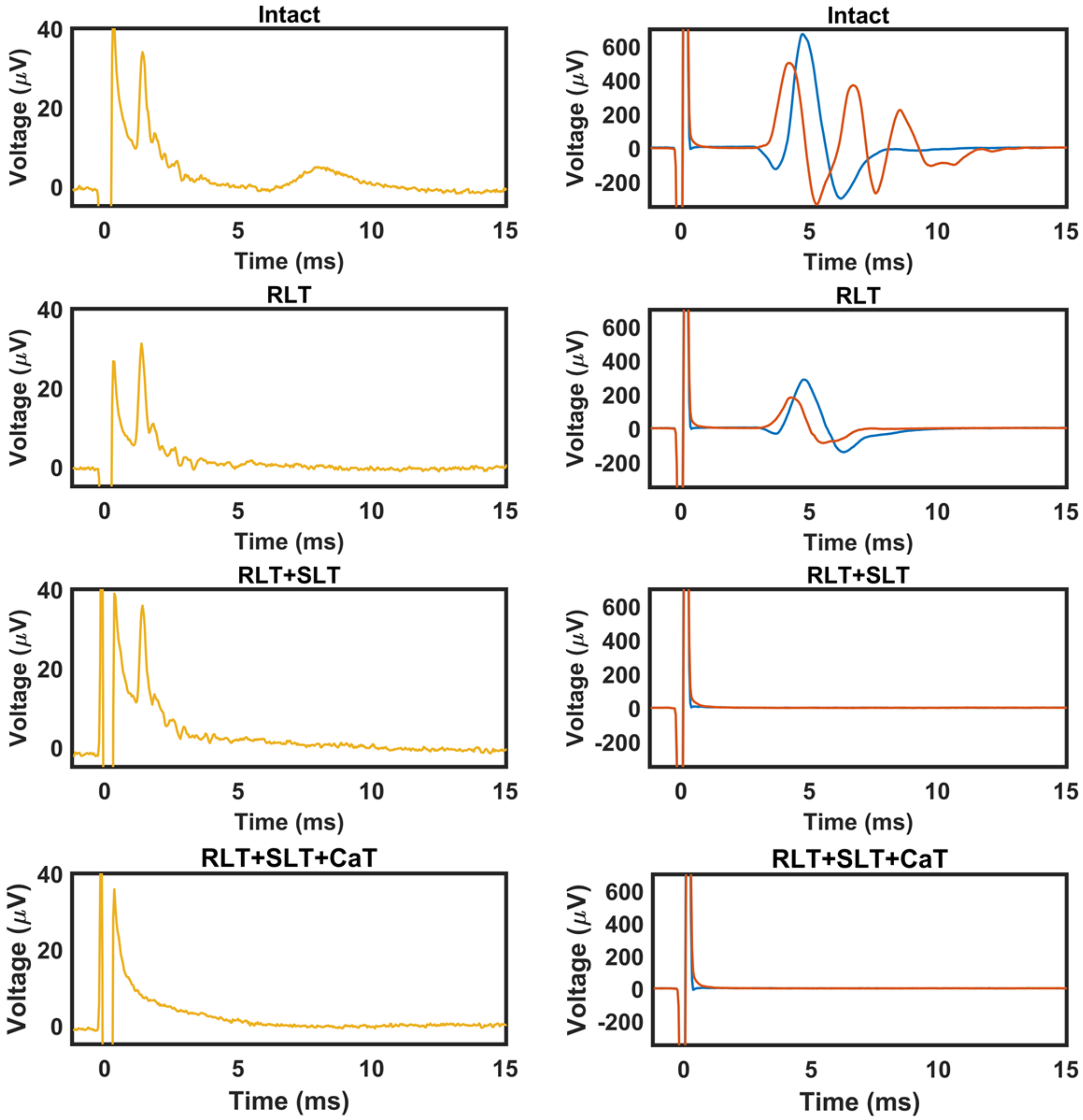

**Supplementary Figure 3**

ENG and EMG traces at every amplitude for intact (no transections and no vecuronium) and vecuronium if applied. Arrows show identified thresholds for ENG fiber types and EMG components. EMG traces without vecuronium were always used to determine EMG component thresholds. ENG traces with vecuronium were used to determine ENG thresholds if vecuronium was applied, otherwise traces without vecuronium were used for determining Aβ- and Aγ-fiber thresholds since the latencies for these signals were always smaller than the fastest short component EMG in any animal. Aδ/B-fiber signals could have the same latency as short or long component EMG signals for any given animal, so Aδ/B-fiber signals were not counted for animals without application of vecuronium. Red arrows in vecuronium traces indicate EMG signals and corresponding ENG artifacts that occurred despite neuromuscular junction block; incomplete block sometimes occurred at the end of stimulation dose response curves (random) as vecuronium was cleared or at higher amplitudes due to vecuronium being a competitive inhibitor and thus can be overcame.

**181205**

Aβ- (300 μA) and Aγ- (1000 μA) fiber signals observed.

Vecuronium incomplete block, possible EMG artifact in ENG signals.

Only short component (2000 μA) EMG observed.

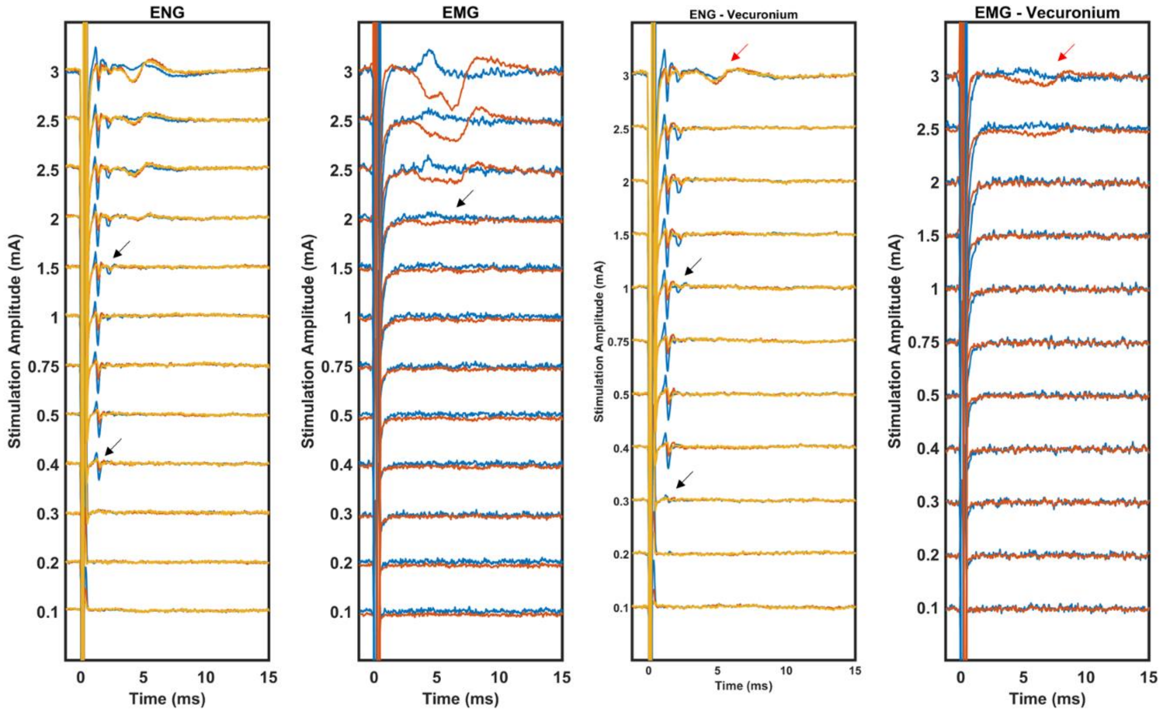

### 181205 ENG Fiber Threshold Zooms

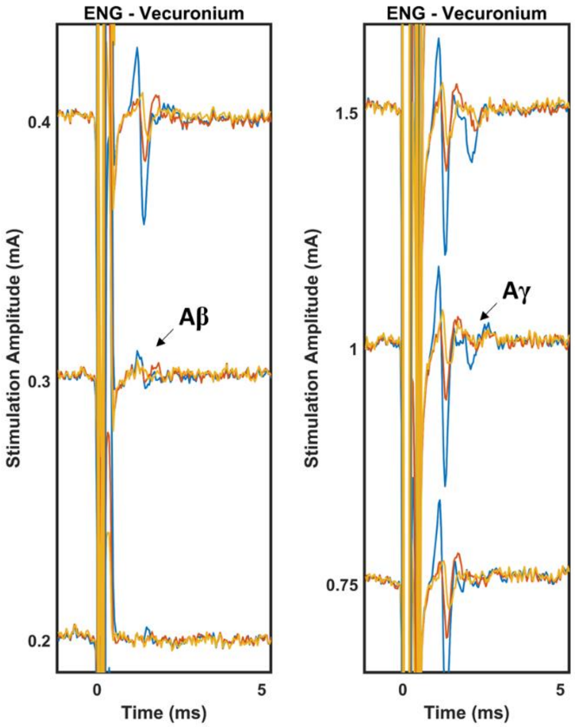

**181211**

Aβ- (200 µA) and Aγ- (1000 µA) fiber signals observed.

Vecuronium not used in this animal, obvious EMG artifact in ENG signals.

Only long component (200 µA) EMG observed.

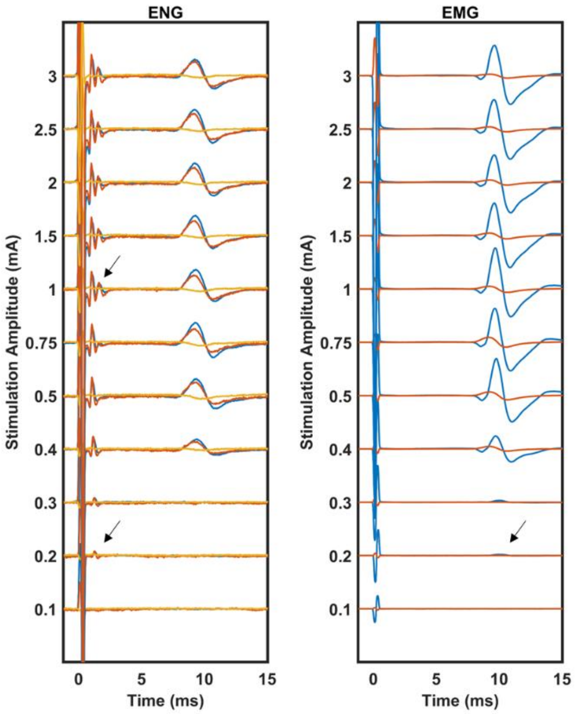

**190109**

Aβ- (300 µA) and Aγ- (1500 µA) fiber signals observed.

Vecuronium incomplete block, no clear EMG artifact or Aδ/B- fiber signals in vecuronium ENG traces.

Only long component (200 µA) EMG observed.

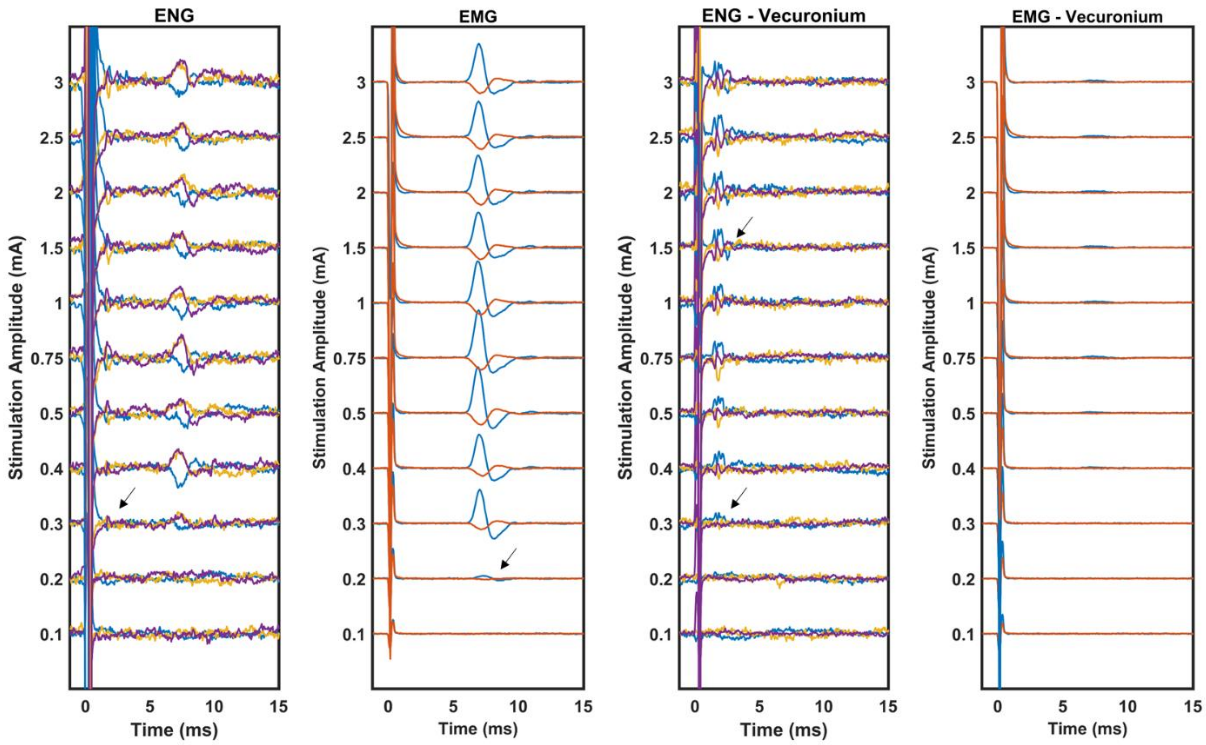

#### 190109 ENG Fiber Threshold Zooms

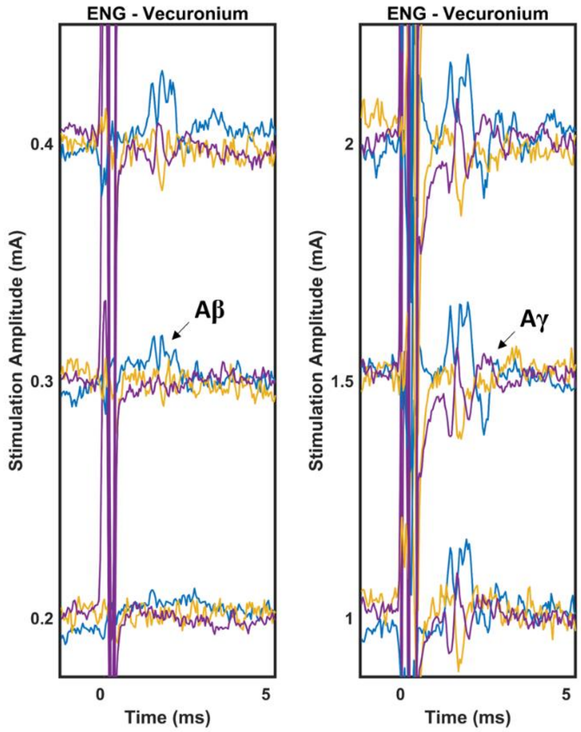

**190122**

Aβ- (300 μA), Aγ- (750 μA), Aδ/B- (2000 μA) fiber signals observed.

Vecuronium complete block.

Short (750 μA) and long component (200 μA) EMG observed.

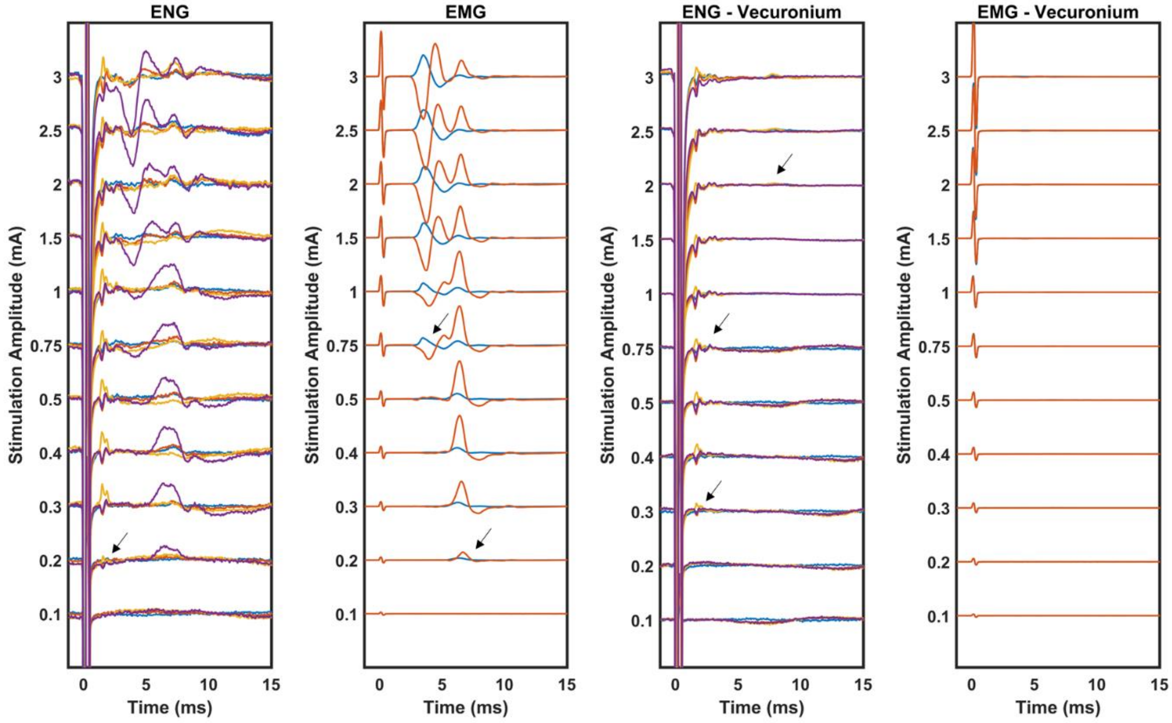

**190122 ENG Fiber Threshold Zooms**

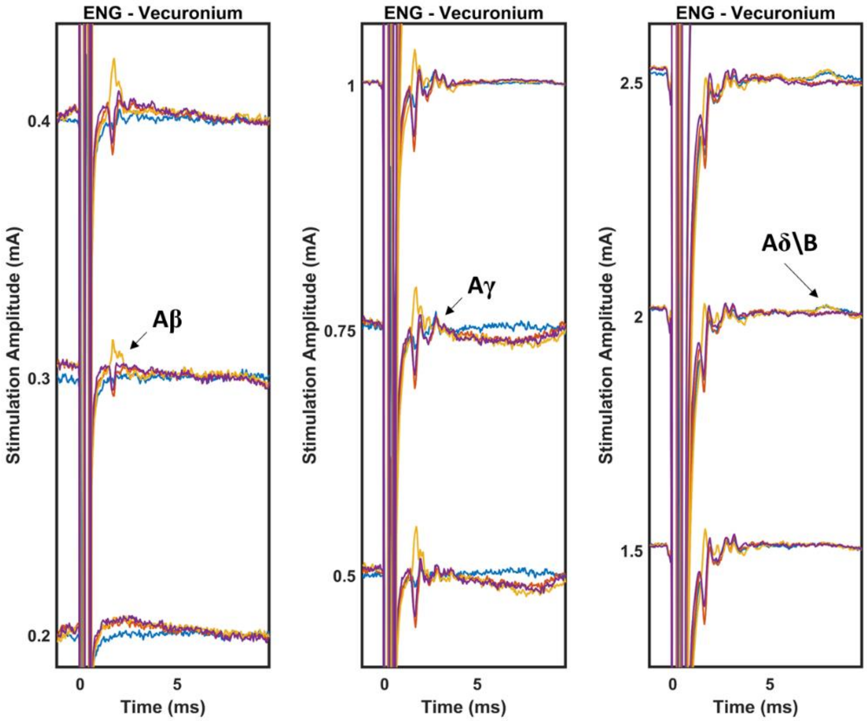

**190122 EMG Component Threshold Zooms**

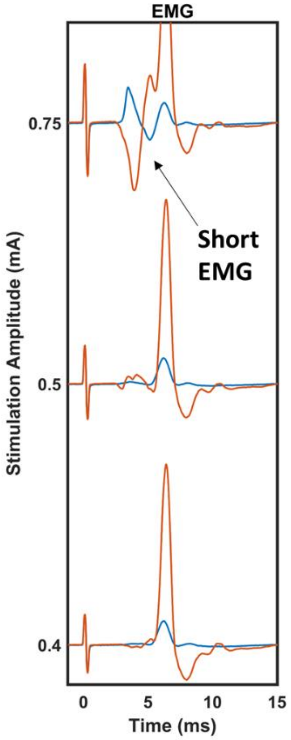

**190123**

Aβ- (100 µA), Aγ- (200 µA), Aδ/B- (1500 µA) fiber signals observed.

Vecuronium incomplete block at some amplitudes. Rational for still identifying Aδ/B- ENG signal is that the EMG signal doesn't match the Aδ/B- signal temporally, and the amplitude of the EMG signal does not correlate with the amplitude of the proposed Aδ/B- signal.

Short (2000 µA) and long component (150 µA) EMG observed.

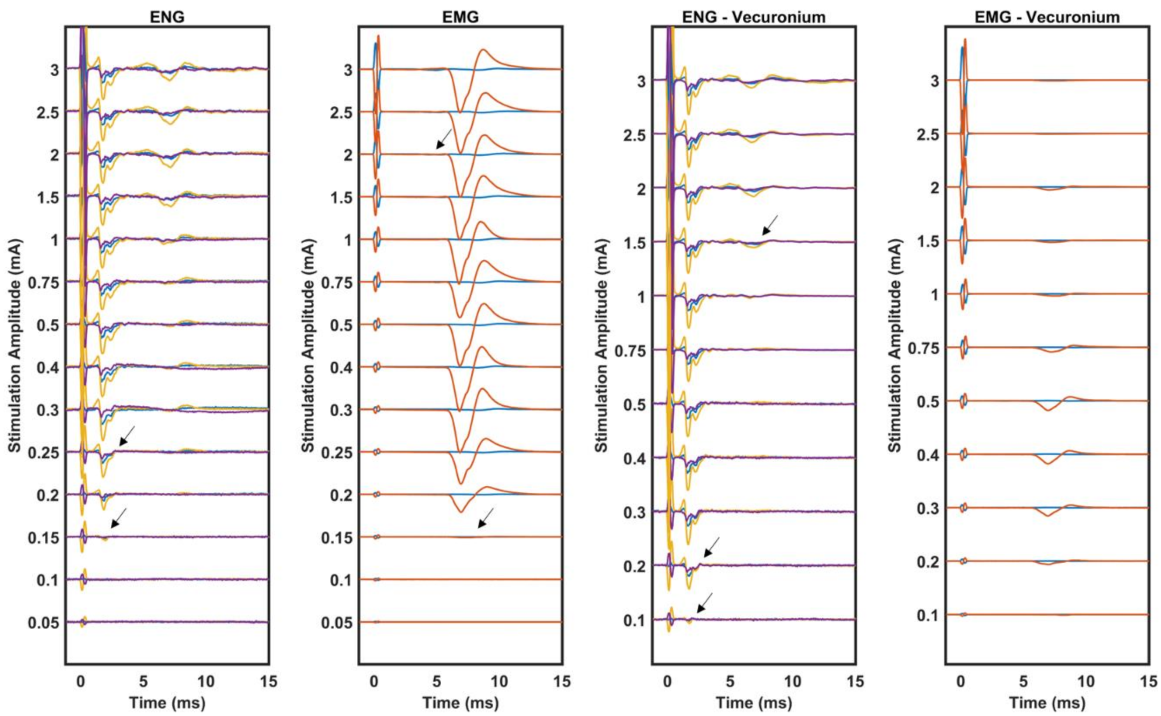

**190123 ENG Fiber Threshold Zooms**

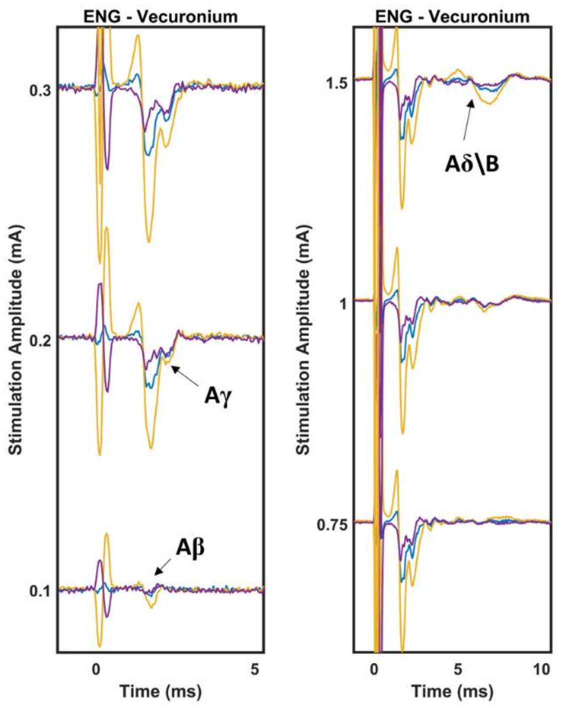

**190123 EMG Component Threshold Zooms**

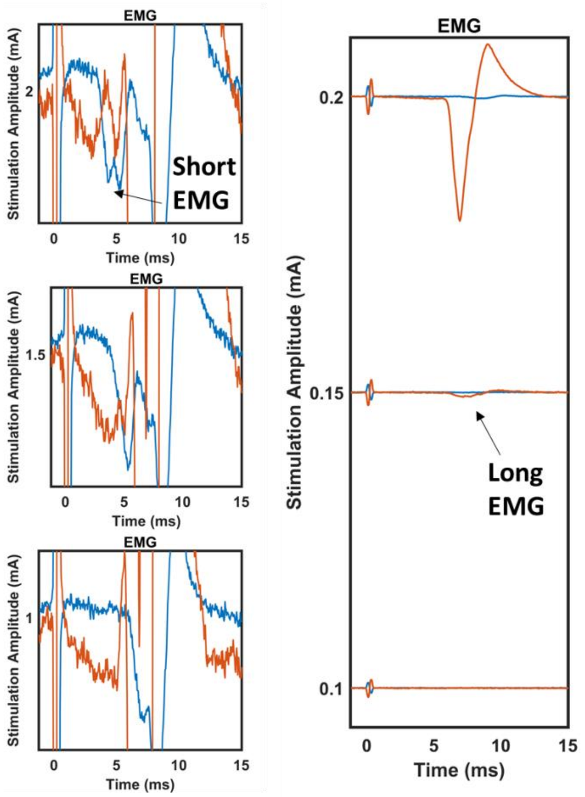

**190129**

Aβ- (200 μA), Aγ- (400 μA), Aδ/B- (1500 μA) fiber signals observed.

Vecuronium incomplete block. Rational for still identifying Aδ/B- ENG signal is that the EMG signal doesn't match the Aδ/B- signal temporally, and the amplitude of the EMG signal does not correlate with the amplitude of the proposed Aδ/B- signal.

Only long component (200 μA) EMG observed.

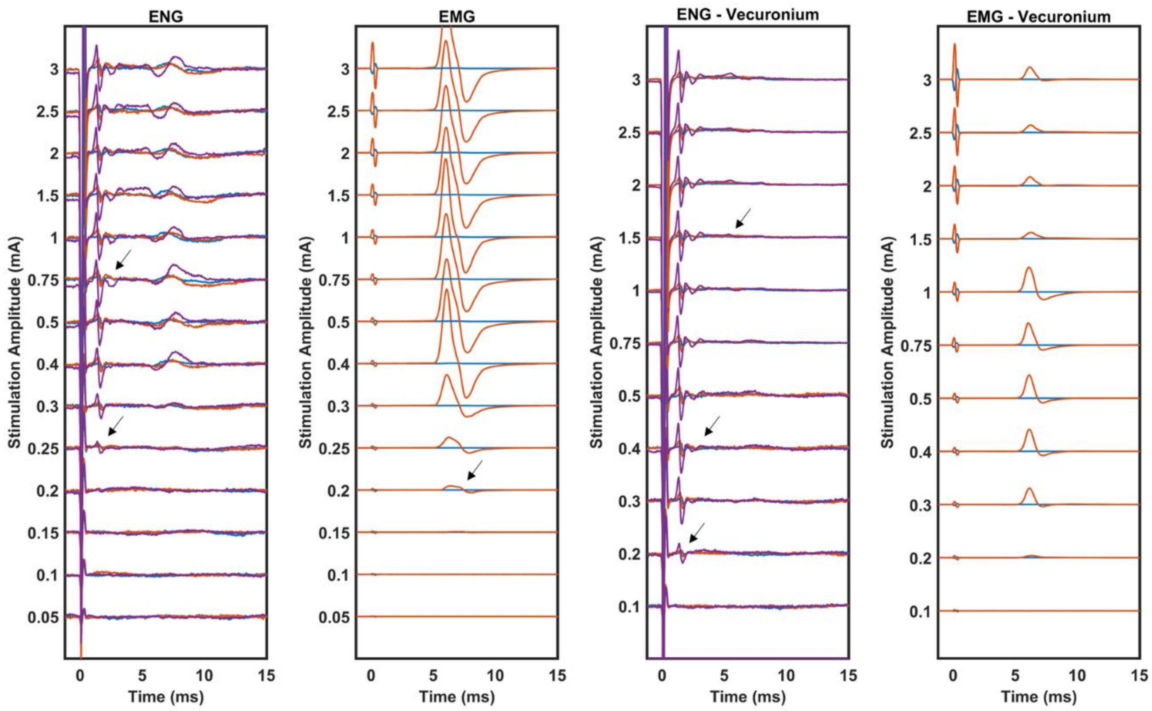

**190129 ENG Fiber Threshold Zooms**

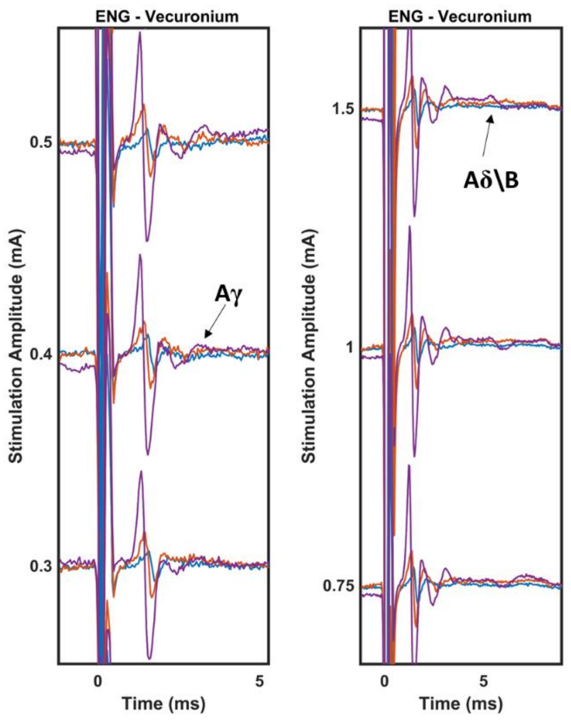

**190227**

Aβ- (200 µA) and Aγ- (300 µA) fiber signals observed.

Vecuronium incomplete block. Block is particularly incomplete at 1500 µA and leads to an EMG response large enough to create an EMG artifact in the ENG trace that could be confused for a compound action potential.

Short (1000 µA) and long component (100 µA) EMG observed.

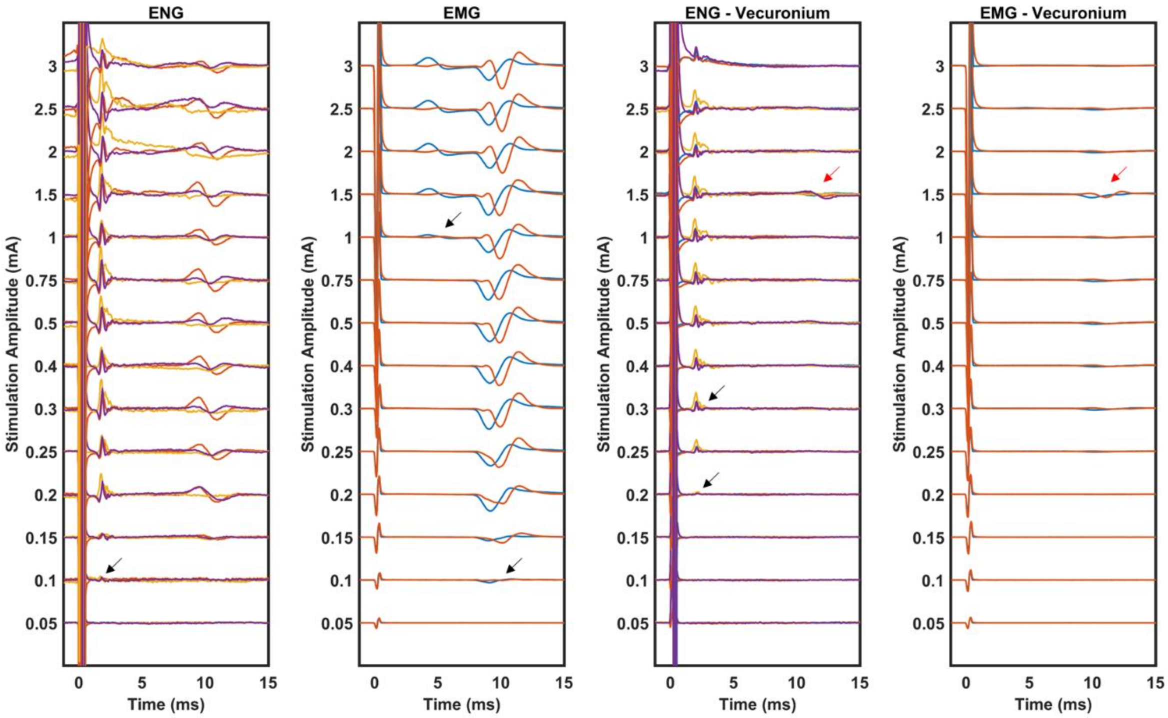

#### 190227 ENG Fiber Threshold Zooms

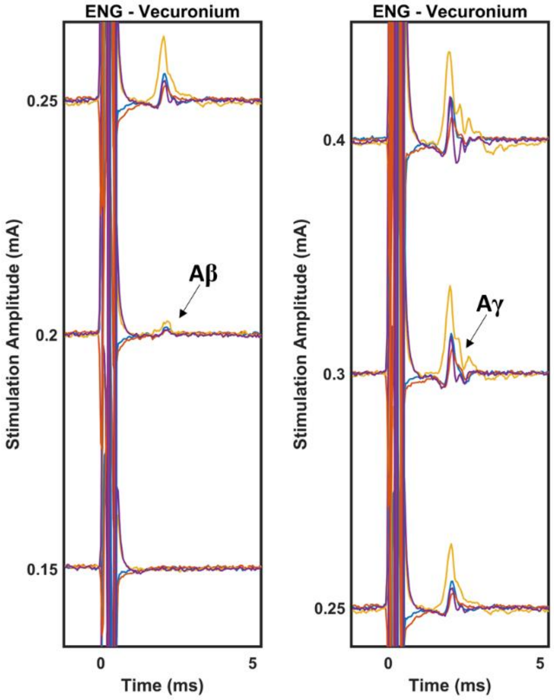

**190403**

Two temporal zooms are presented. First, 0 to 15 seconds to match the rest of the animals. Second, 0 to 30 seconds to show the possible C-fiber, which was not observed in any of the other animals (hence not shown for any other animals).

Aβ- (500 µA) and Aγ- (1500 µA) fiber signals observed. Possible C-fiber at 2000 µA.

Vecuronium was not used in this animal, though branch transections show that short and long EMG components can be completely removed leaving observable Aβ-, Aγ-, and C-fiber signals, which are then removed by transection of the main vagus trunk.

Short (1500 µA) and long component (750 µA) EMG observed.

Some rational behind hesitation on C-fiber identification. No vecuronium in this animal, only branch transections; C-fiber signal could be caused by some unidentified motor pathway. Thresholds for all signals were generally higher in this animal than the rest of the cohort, which should mean any C-fiber should have a higher threshold than any other animal, yet this is the only animal with a C-fiber signal. No observation of Aδ/B- signals, which should have lower thresholds than C-fibers.

From our perspective, the only explanation is that the ENG LIFEs were placed directly into fascicles containing C-fibers and were far away from fascicles containing Aδ or B fibers. Hence the hesitation.

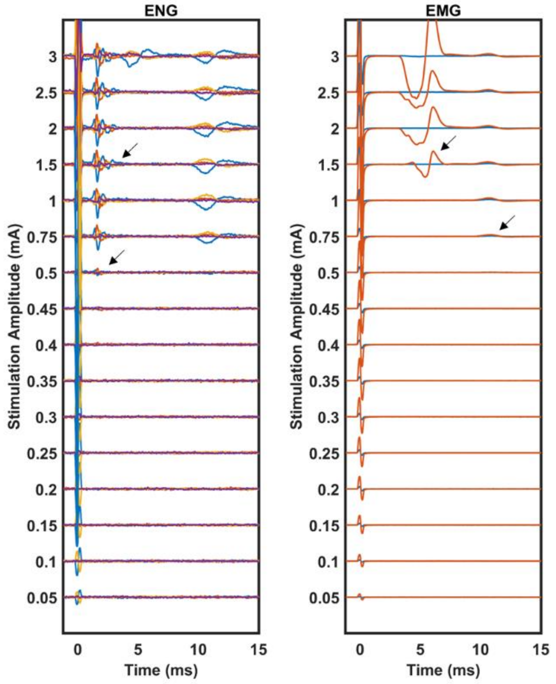

**190403 Temporal Expansion for Possible C-fiber**

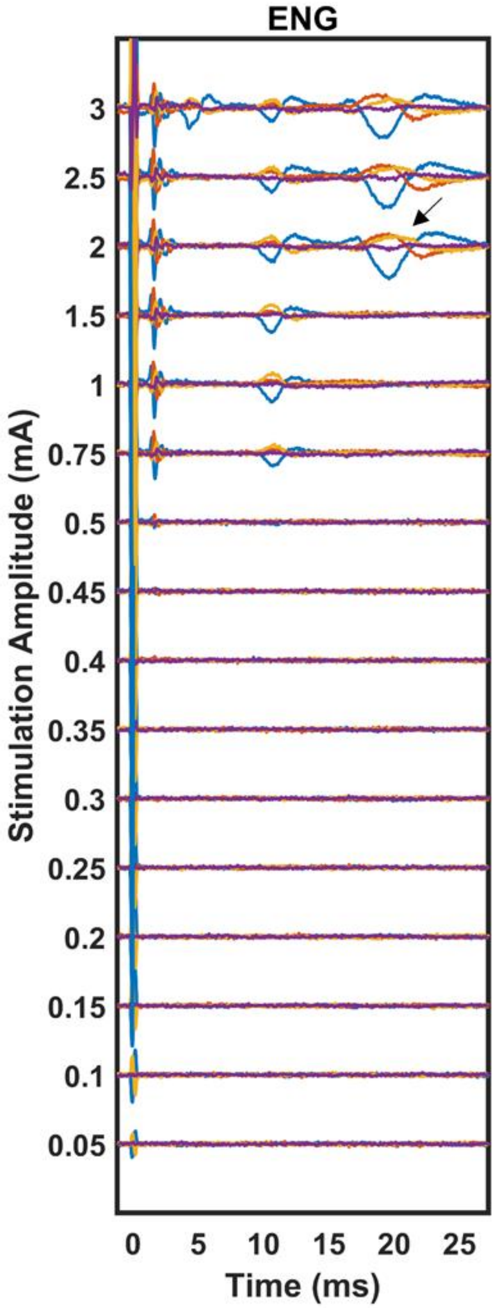

#### 190403 ENG Fiber Threshold Zooms

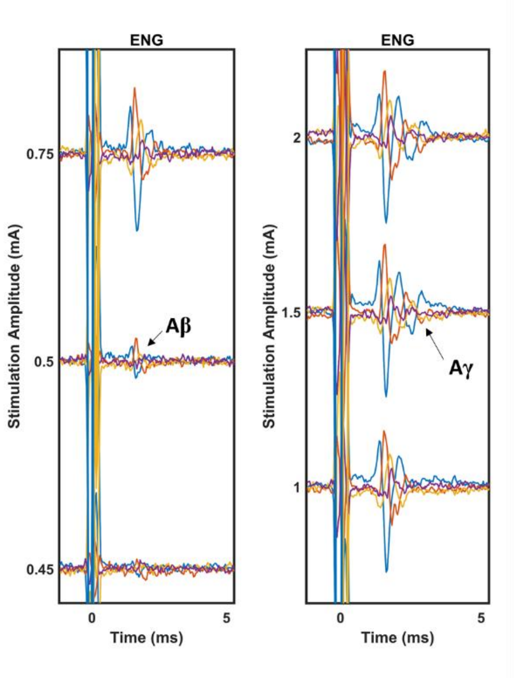

### 190403 EMG Component Threshold Zooms

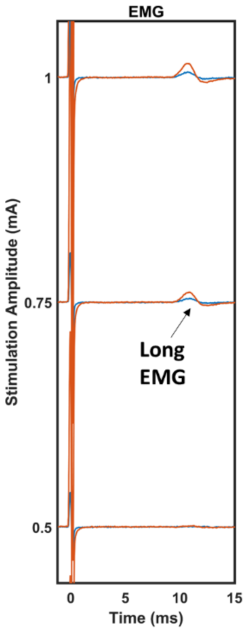

**190410**

Aβ- (250 µA) and Aγ- (450 µA) fiber signals observed.

Vecuronium was not used in this animal.

Short (2500 µA) and long component (350 µA) EMG observed.

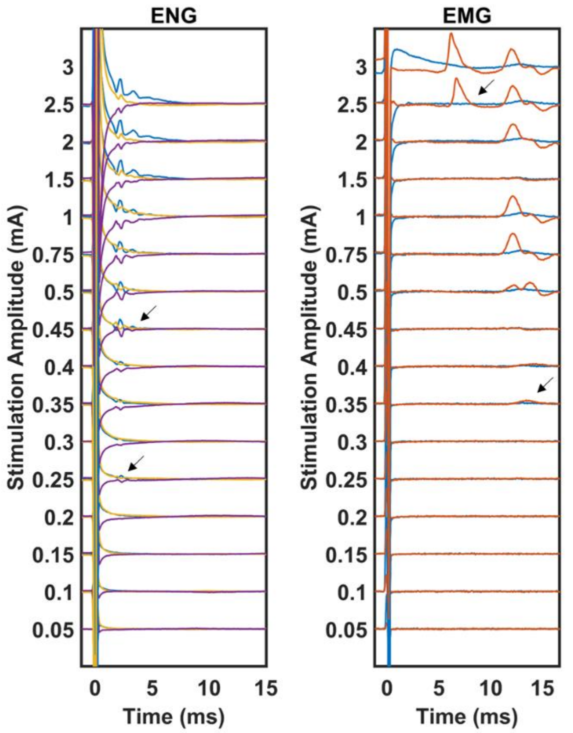

**190410 ENG Fiber Threshold Zooms**

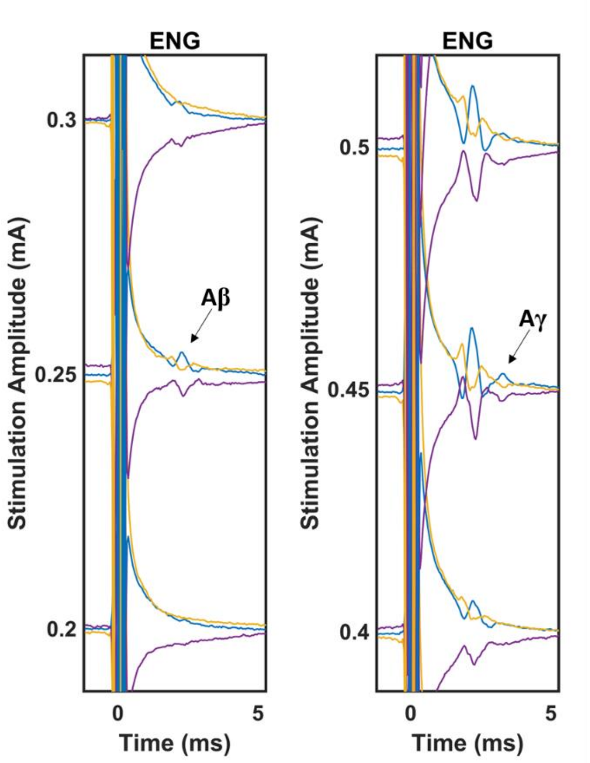

**190410 EMG Component Threshold Zooms**

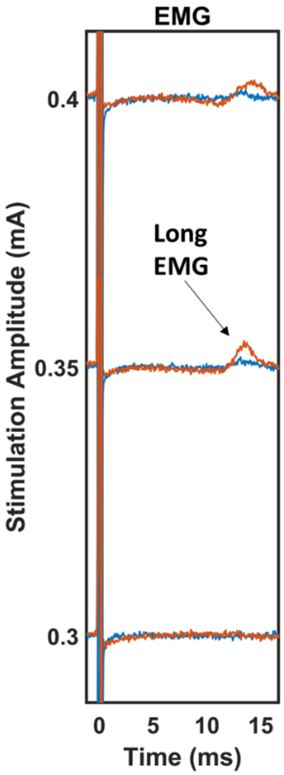

**190501**

Only Aβ- (200 µA) fiber signals observed.

Vecuronium complete block.

Short (1500 µA) and long component (350 µA) EMG observed.

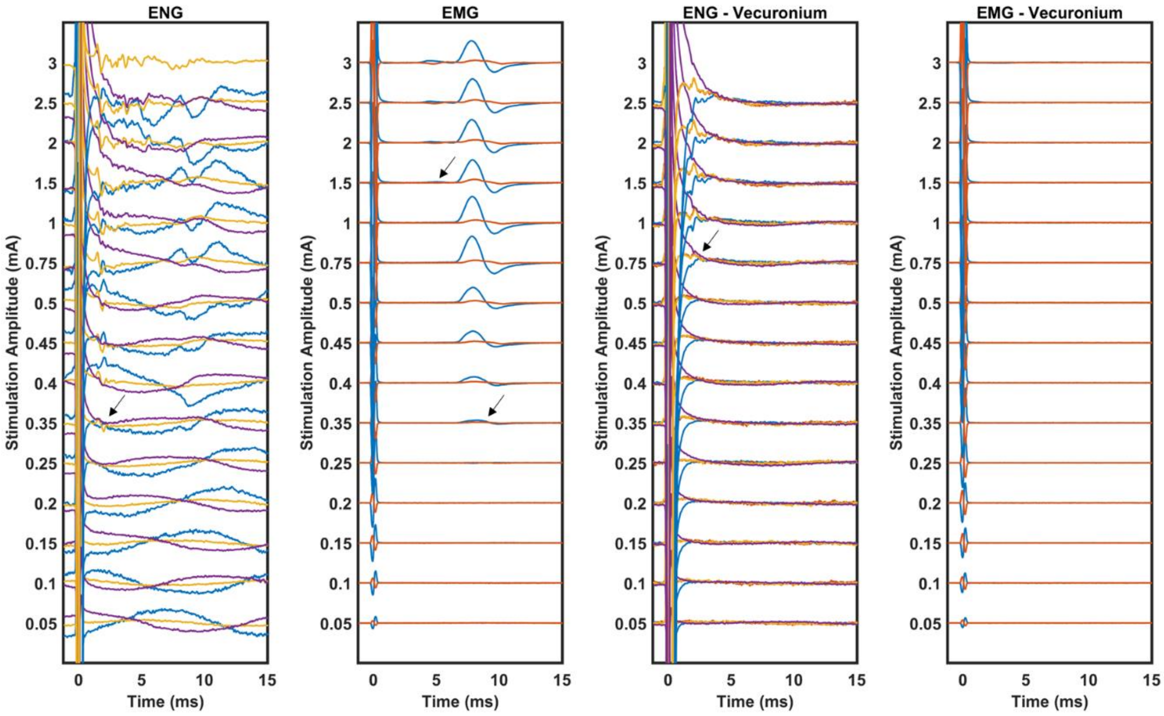

#### 190501 ENG Fiber Threshold Zooms

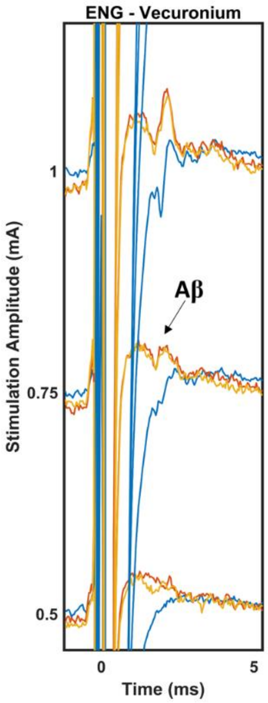

**190501 EMG Component Threshold Zooms**

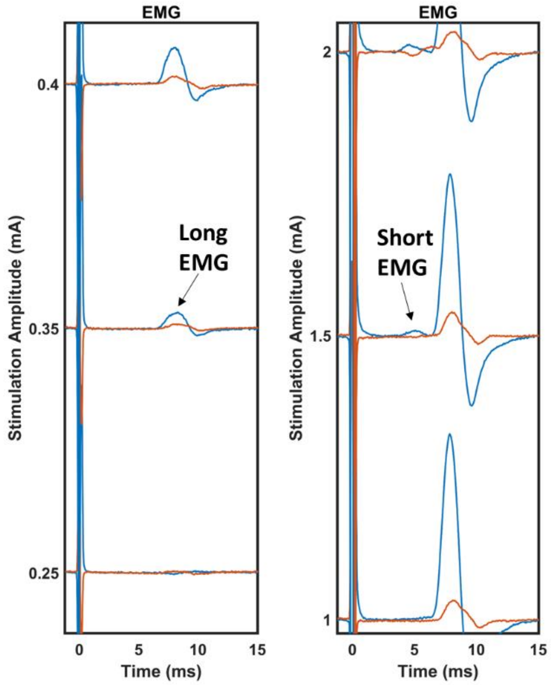

**190522**

Only Aβ- (750 µA) fiber signals observed.

Vecuronium was not used in this animal, though both branch transections were performed.

Short (500 µA) and long component (400 µA) EMG observed.

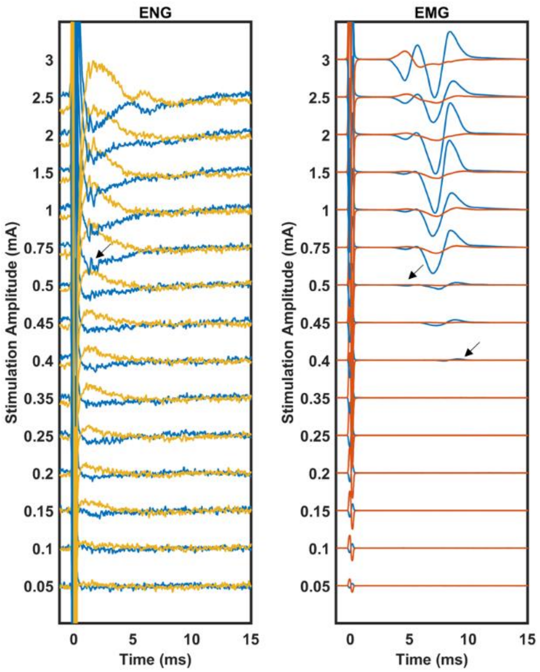

**190522 EMG Component Threshold Zooms**

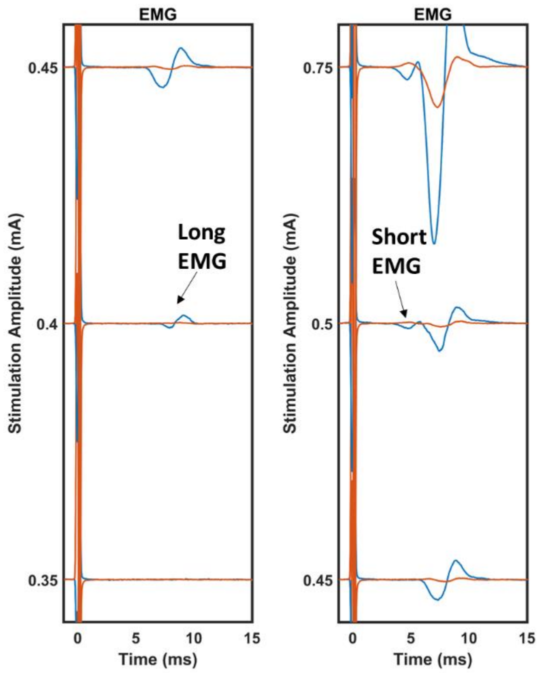

**190529**

Only Aβ- (400 µA) fiber signals observed.

Vecuronium was not used in this animal, though both branch transections were performed.

Short (1500 µA) and long component (350 µA) EMG observed.

Subtracted channel 4 from channel 2 for this figure.

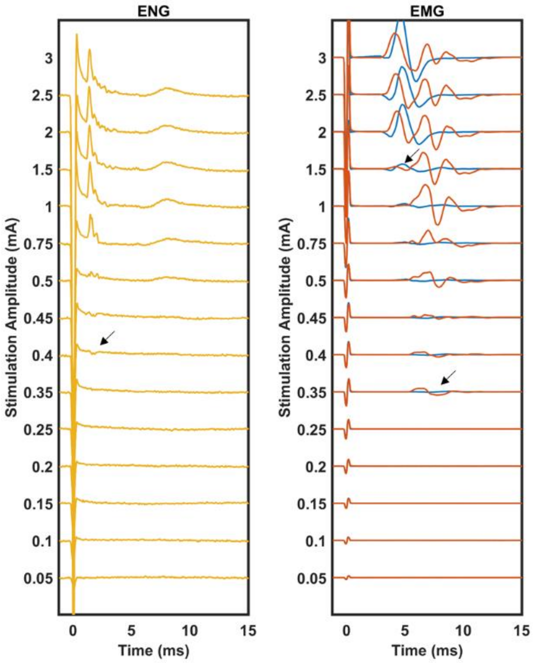

**190529 ENG Fiber Threshold Zooms**

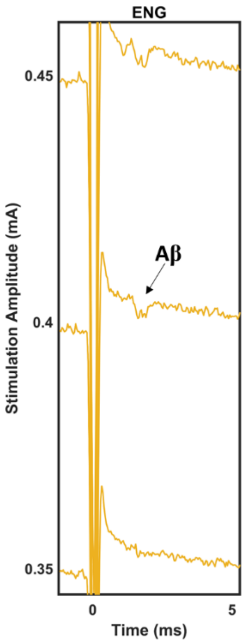

**Supplementary Figure 4**

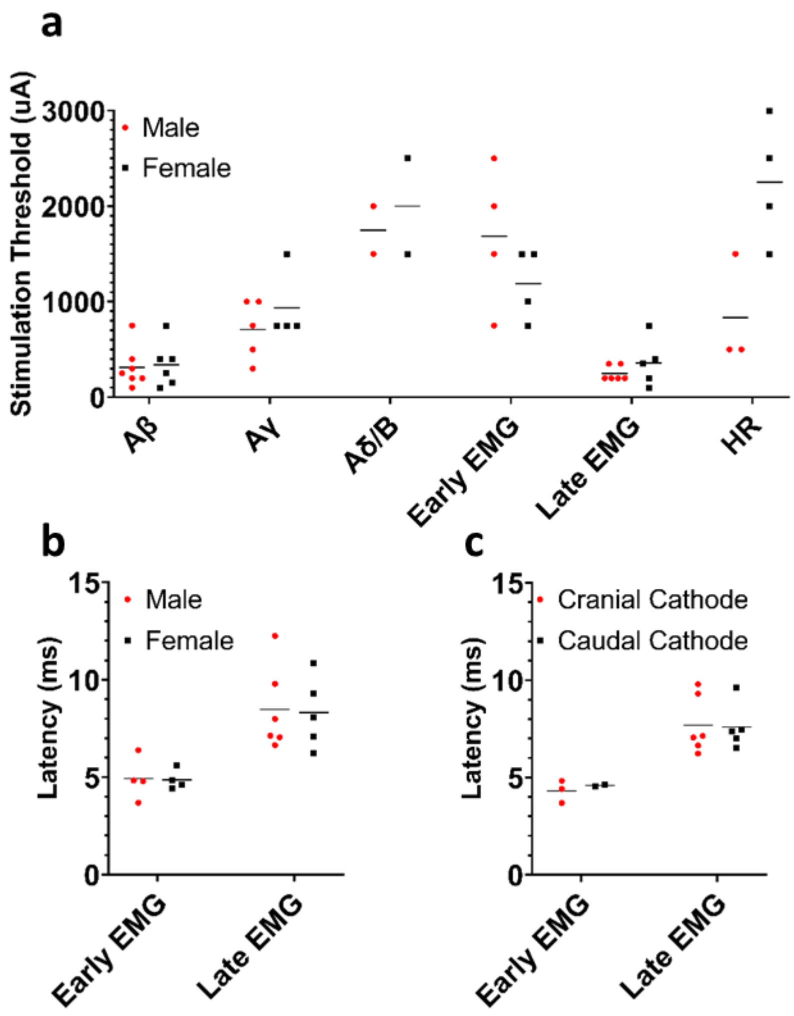


**Supplementary Figure 4: Additional Comparisons Between Male and Female Pigs, as well as Cathode and Anode Configurations. a)** Thresholds for ENG, EMG, and HR responses comparing male and female pig experiments. **b)** Post-stimulus latencies for EMG components comparing male and female pig experiments. **c)** Post-stimulus latencies for EMG components comparing anode and cathode configurations. Note that only animals where responses to both cathode cranial and cathode caudal configurations were recorded are plotted for panel c.

**Supplementary Figure 5**

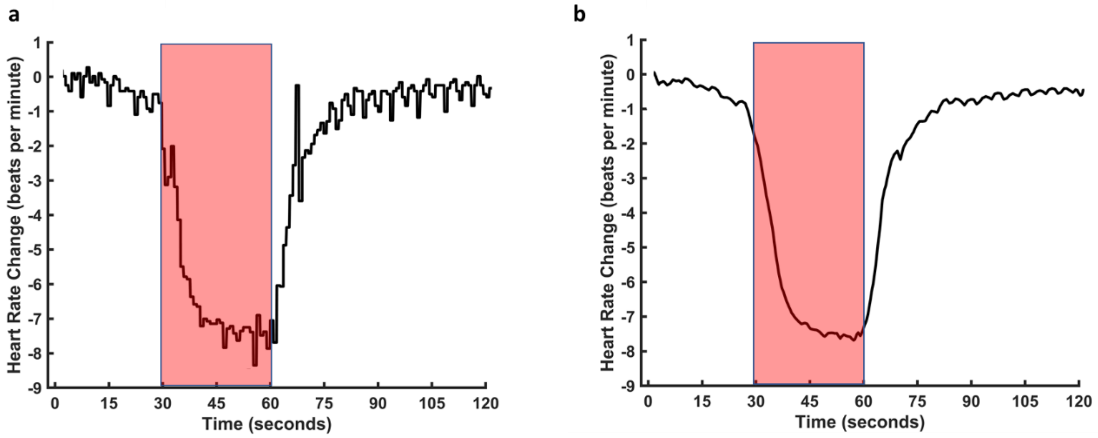


**Supplementary Figure 5: Example of Vagus Nerve Stimulation Evoked Heart Rate Decrease. a)** Raw heart rate trace with 30 seconds of beginning baseline, 30 seconds of stimulation (red transparent box), and 60 seconds of ending baseline. **b)** Same data, but smoothed with 1000 point Matlab smooth function. Smoothed version was used to calculate changes in all animals. The median of the 30 seconds preceding stimulation was used as the baseline value. The minimum value of the 30 seconds during stimulation was subtracted from the baseline value to calculate the change in heart rate due to stimulation.

**Supplementary Figure 6**

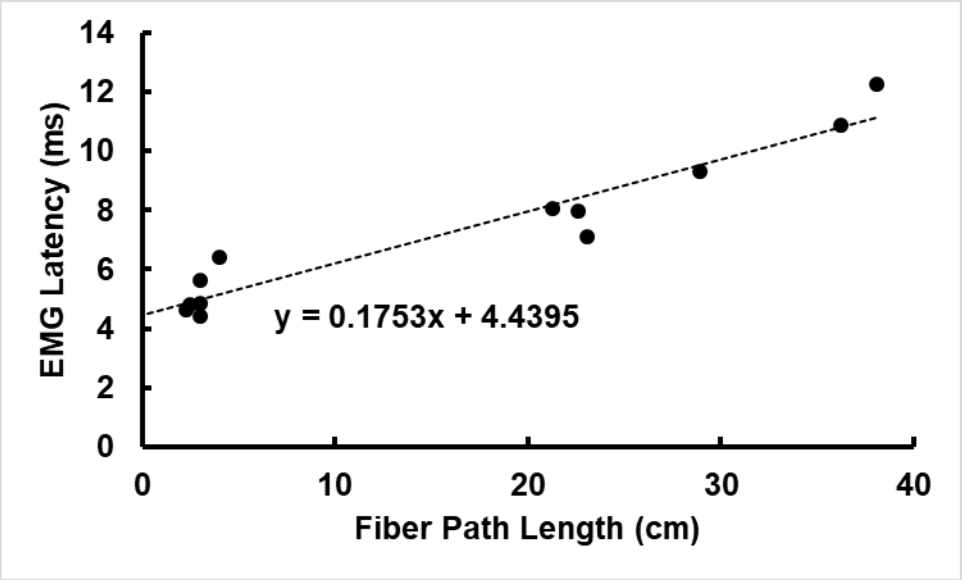


**Supplementary Figure 6: Determination of Synaptic Delay at Neuromuscular Junction of Pig Vagus Nerve Somatic Branches and Innervated Neck Muscles.** The length of nerve fiber that an action potential would have been expected to travel (Fiber Path Length, x-axis) from the stimulation electrode to the cricothyroid and cricoarytenoid muscles via the superior and recurrent laryngeal branches was correlated to the corresponding EMG post-stimulus response latency (EMG Latency, y-axis) generated by each path as determined by branch transection data. The path length for the recurrent laryngeal branch was determined as the distance between the cranial stimulation contact (cathode for all cases plotted) and the recurrent laryngeal branching point added to the distance between the recurrent laryngeal branching point to insertion into the cricothyroid muscle (see Supplementary Table 1, row “Cranial E Recurrent Path Length”). The path length for the superior laryngeal branch was determined as the distance between the superior laryngeal branching point from the vagus trunk to insertion into the cricothyroid muscle (see Supplementary Table 1, row “Superior Laryngeal Length”); the distance between the cranial stimulation contact was not factored into the path length for the superior laryngeal since the fibers do not pass under the electrode cuff. The y-intercept of the trendline for this linear relation ($R^2 = 0.92$) should be representative of the synaptic delay at the neuromuscular junction since a fiber path length of 0 cm would be expected to still have an EMG post-stimulus latency of approximately 4.4 ms.
